# Strategies to assure optimal trade-offs among competing objectives for genetic improvement of soybean

**DOI:** 10.1101/2021.02.19.431938

**Authors:** Vishnu Ramasubramanian, William Beavis

## Abstract

Plant breeding is a decision making discipline based on understanding project objectives. Genetic improvement projects can have two competing objectives: maximize rate of genetic improvement and minimize loss of useful genetic variance. For commercial plant breeders competition in the marketplace forces greater emphasis on maximizing immediate genetic improvements. In contrast public plant breeders have an opportunity, perhaps an obligation, to place greater emphasis on minimizing loss of useful genetic variance while realizing genetic improvements. Considerable research indicates that short term genetic gains from Genomic Selection (GS) are much greater than Phenotypic Selection (PS), while PS provides better long term genetic gains because PS retains useful genetic diversity during the early cycles of selection. With limited resources must a soybean breeder choose between the two extreme responses provided by GS or PS? Or is it possible to develop novel breeding strategies that will provide a desirable compromise between the competing objectives? To address these questions, we decomposed breeding strategies into decisions about selection methods, mating designs and whether the breeding population should be organized as family islands. For breeding populations organized into islands decisions about possible migration rules among family islands were included. From among 60 possible strategies, genetic improvement is maximized for the first five to ten cycles using GS, a hub network mating design in breeding populations organized as fully connected family islands and migration rules allowing exchange of two lines among islands every other cycle of selection. If the objectives are to maximize both short-term and long-term gains, then the best compromise strategy is similar except a genomic mating design, instead of a hub networked mating design, is used. This strategy also resulted in realizing the greatest proportion of genetic potential of the founder populations. Weighted genomic selection applied to both non-isolated and island populations also resulted in realization of the greatest proportion of genetic potential of the founders, but required more cycles than the best compromise strategy.

## 1. Background

Historical responses to selection of commodity crops has been enabled by decreasing the number of years between cycles of recurrent selection, by increasing the number of replicable genotypes (selection intensity) and by increasing the number of field trials (heritability on an entry mean basis). In other words, genotypic improvements from responses to selection in commodity crops over the last 50 years (Specht et al. 2014) required monetary investments that became part of the exponential rise in seed costs during the same time (Byrum et al. 2017; USDA-ERS 2020). Since the emergence and adoption of Genomic Selection (GS), it has been possible to increase the numbers of genotypes that are evaluated, i.e., selection intensity, without significant increases in numbers of field plots (Bernardo 2007, 2008; Asoro et al. 2011; Heslot et al. 2012; Nakaya and Isobe 2012; Emily and Bernardo 2013; Crossa et al. 2014; Beyene et al. 2015; Bassi et al. 2016; Marulanda et al. 2016; Jonas and de Koning 2016; Hickey et al. 2017; Goiffon et al. 2017).

In a companion manuscript we reported an investigation of various factors on response metrics to recurrent selection of soybean lines derived from founders of the SoyNAM population (Ramasubramanian and Beavis 2020). The combinatorial set of factors consisted of phenotypic selection (PS) and four commonly used GS methods, training sets (TS), selection intensity (SI), number of QTL (nQTL), and heritability (H) on an entry mean basis. While interactions among all factors affected all response metrics, only the impacts of GS methods, SI and TS are factors that plant breeders can control. All GS methods provided greater responses than PS for at least five cycles, but PS provided better responses to selection as response from GS methods reached a limit. These results are consistent with reports by Goddard (2009), Jannink (2010) and Liu et al. (2015) demonstrated that the full genotypic potential of the founders is eliminated more quickly with GS than PS. In terms of factors that a soybean breeder can control, we found that SI’s of 1.75 and use of Ridge Regression Genomic Prediction (RRGP) models updated every cycle with training data from all prior cycles of selection provided rapid response in the early cycles of selection and retention of genetic diversity for continued response to selection in later cycles, but we noted that further improvements might be made if the populations were organized into islands and mating designs other than the hub network were employed (Ramasubramanian and Beavis 2020). Herein we investigate strategies that soybean breeders can employ to find optimal trade-offs between maximizing genetic gain from selection and retaining useful genetic diversity.

The challenge of realizing genetic gains from selection and retaining useful genetic diversity in closed populations has been of interest since it was demonstrated that there are theoretical limits for response to selection in closed populations (Hill and Robertson 1968; Bulmer 1971). Trade-offs among objectives don’t prohibit finding optima as long as optimality is defined as a compromise among competing objective functions (Deb 2003; Konak et al. 2006; Shoval et al. 2012; Sheftel et al. 2013; Saeki et al. 2014).

Before the development of GS, quantitative geneticists working on domestic animal systems utilized mathematical programming modeling and operations research (OR) approaches to find near-optimal solutions to the challenge of assuring genetic gain and minimizing inbreeding per cycle of selection (Wray and Goddard 1994). The first publication using OR approaches to address multiple objectives in plant breeding was applied to selection of multiple traits (Johnson et al. 1988). Generally OR approaches involve three activities: 1) define objectives using measurable metrics, 2) translate the objectives into mathematical programming models consisting of objective functions, decision variables and constraints, 3) find an algorithm that will provide values for the decision variables resulting in optimal solutions to the mathematical programming model (Rardin 2017).

If the genetic improvement project wants to assure genetic gain and retain useful genetic diversity then there are two competing objectives for which a trade-off needs to be optimized. This represents an example of a multi-objective optimization (MOO) problem (Deb 2003, 2011; Rardin 2017). After translating each of the objectives into an objective function, there are several strategies for finding the optimal solution (Deb 2003). The two most commonly used strategies are known as the ε -constraint and the weighted sum. The ε -constraint method consists of identifying one of the objectives, e.g., maximize genetic gain, and translate other objectives, such as minimize inbreeding, into decision variables that can be constrained in a linear, integer or quadratic mathematical programming model (Haimes 1971). In other words, translate the MOO mathematical model into a Single Objective Optimization (SOO) model for which there exist computational algorithms capable of finding the optimum solution (Frank and Wolfe 1956; McCarl et al. 1977; Lazimy 1982). Framing the ε -constraint method requires definition of metrics for genetic diversity or inbreeding. In animal breeding this method became known as Optimum Contribution Selection (OCS: Wray and Goddard 1994; Brisbane and Gibson 1995; Meuwissen 1997; Grundy et al. 1998; Meuwissen et al. 2001). Subsequent to development of GS, OCS was modified to maximize Genomic Estimated Breeding Values (GEBVs) and the realized relationship matrix was used to constrain inbreeding in what became known as Genomic OCS (GOCS) (Sonesson et al. 2010; Schierenbeck et al. 2011; Woolliams et al. 2015).

The second well-established approach to a MOO challenge is known as the weighted sum method. The weighted sum method assigns weights, ω_i_ ∈ [0, 1] and ∑ ω_i_ =1, to each of the ‘i’ objective functions and an algorithm is employed to find the values for the decision variables that minimize all objective functions simultaneously (Zadeh 1963). Breeders will recognize the weighted sum method as a selection index composed of weighted parameters for genetic gain and inbreeding, or equivalently genetic diversity. If genomic information is available, GEBV’s can be used to maximize genetic gain and the realized relationship matrix can be used to minimize inbreeding resulting in a genomic selection index (GSI) that can be calculated for all genotypes. (Carvalheiro et al. 2010; Clark et al. 2013).

Both ε-constraint and weighted sum methods are referred to as preference methods (Deb 2003) where the constraints or relative weights have been predetermined. For defined preferences there exist exact optimization algorithms if Karush-Kuhn-Tucker (KKT) conditions are met (Karush 1939; Kuhn and Tucker 1951). An exact optimization solution guarantees that no other feasible solution will be a better solution for the specified set of constraints or weights. Unfortunately, it is difficult to predetermine these values because they require forecasting the relative economic values of genetic gains and retention of useful genetic diversity. For commercial plant breeding projects competition in the marketplace will force much greater emphasis on maximizing genetic gains than retaining genetic diversity. In contrast public soybean breeders have an opportunity, perhaps an obligation, to retain useful genetic diversity while realizing genetic gains for quantitative traits of agronomic importance. Because each plant breeding project has unique relative trade-offs, evolutionary algorithms (EAs) have been adopted to provide multiple solutions on an efficient (Pareto) frontier of solutions to competing objectives (Deb 2003, 2011; Konak et al. 2006). Decision makers then decide which of the solutions have the appropriate relative emphasis on the competing objectives.

Genetic algorithms (GAs) are a class of EAs that are based on recurrent selection of breeding populations and were developed to find computational solutions to large combinatorial problems (Goldberg 1989; Luque 2011). In a canonical GA, selected solutions are pooled together into a set of solutions. Subsequently the individual solutions are randomly sampled for pairwise “matings” to create a new set of solutions for evaluation and selection. The algorithm is iterated until there are no improvements in the sets of solutions. Computational analogs of mutation or recombination, are utilized to move from local optima to global optima. A subclass of GAs, known as parallel GAs maintain structure among subsets of individual solutions and enable the subsets to independently find different solutions for different domains (Luque 2011). The parallel GA system is analogous to the concept of genetic subpopulations (Falconer and Mackay Figure 3.2, 1996). Island Model GAs allow for exchange of solutions among subpopulations. Island model GAs (IMGAs) are distinct from canonical GAs in terms of properties and behavior because evolution happens locally, within island, and globally, among islands. Island model parameters consist of number of islands, island size, selection pressure within each islands, numbers of migrants, migration frequency, connectedness or topology of islands and emigration and immigration policies among islands (Whitley 1999; Skolicki 2007 a, b).

Rather than investigate the trade-off between objective functions, Jannink (2010) proposed that it would be possible to retain useful genetic diversity in GS by weighting low frequency alleles with favorable estimated genetic effects. Simulations with Weighted Genomic Selection (WGS) resulted in greater responses across 24 selection cycles of recurrent selection than unweighted GS, using RRBLUP values, for both low and high heritability traits. However, the initial rates of response using WGS were less than responses from application of PS and less than GS. The response using WGS was better than response from PS after twenty cycles of selection, but the responses relative to GS depended on the number of simulated QTL and heritability. Decay of LD between marker and QTL is one of the factors that can slow responses using GS relative to PS (Hickey et al. 2014; Xavier et al. 2016), although decay of LD did not contribute to responses in the initial cycles using WGS. The rate of inbreeding per cycle is also greater with GS than with PS, whereas it is similar to PS when WGS is applied. The rate of fixation of favorable alleles is lower for WGS than GS resulting in larger numbers of cycles of genetic improvement before response to selection reaches a limit (Jannink 2010). Efforts to balance the response in early cycles and later cycles have included addition of parameters to WGS (Sun and Van Raden, 2014) and dynamic weighting of rare alleles depending on the time horizon for the breeding program (Liu et al., 2015). Low frequency favorable alleles are given greater weights, drawn from a Beta distribution, in initial cycles, and the weights tend towards unity as the number of cycles of selection approaches a predefined time horizon. This shifts the balance towards retaining greater genetic variance in earlier cycles.

The applications of GS, GOCS, GSI and WGS assume that selected individuals will be randomly mated. Typically, plant breeders do not randomly mate selected genotypes, rather most use selected genotypes that exhibit the most desirable selection metrics, e.g., GEBVs, to serve as “hub” parents in networked crossing designs (Guo et al. 2013; Guo et al. 2014). Such Hub-Network (HN) mating designs (MD’s) apply greater weights to genetic contributions from hub genotypes resulting in amplified loss of genetic diversity relative to random mating by reducing the effective population size.

As soybean breeders have become aware of the potential impacts due to loss of genetic diversity from use of GS, they have used various *ad hoc* methods to avoid crosses between related genotypes (Diers, Graef, Lorenz, Cianzio, Singh, personal communications). After quantitative geneticists working on animal breeding systems demonstrated that it is possible to use the GSI strategy with an EA to identify optimal pairs of mates (Kinghorn 2011; Pryce et al. 2012; Woolliams et al. 2015), plant quantitative geneticists developed and investigated various versions of GSI and GOCS for plant breeding (Akdemir and Sanchez 2016; Lin et al. 2017; Cowling et al. 2017; Beukelaer et al. 2017; Gorjanc et al. 2018; Allier et al. 2019). Notice that the computational demand to find the optimum on the non-decreasing efficiency frontier created by all possible constraint values or relative weights in all *NC* _2_ mating pairs is particularly well suited for application of EA’s. Also, it should be noted that Akdemir and Sanchez (2016) referred to their implementation of GOCS as efficient GS. Last, we note that optimal mate selection has been referred to as optimal cross selection in plant breeding applications (Gorjanc et al. 2018; Allier et al. 2019), unfortunately with the same acronym as OCS. To distinguish optimal cross selection from optimal contribution selection, we do not use an acronym for optimal cross selection.

In addition to evaluating traditional PS, GS, and GOCS, Akdemir and Sanchez (2016) proposed and evaluated a novel mathematical programming model, referred to as genomic mating (GM). They formulated the problem as minimizing a linear function of inbreeding plus a negative risk function for the realized relationship matrix of N_p_ possible parents. Inbreeding is a function of the expected genetic diversity among N_c_ progeny from the N_p_ parents and is weighted by a parameter that controls allelic diversity among all N_p_ parents. Risk is determined for each cross as the sum of the expected breeding values of the progeny plus the expected standard deviations of marker loci weighted by a parameter that controls allelic heterozygosity of the relative contributions of the marker loci to the GEBVs. Thus, risk is similar to the usefulness criterion defined by Schnell (1983 as cited in Melchinger et al. 1988) of a selected proportion of the population and the weighting parameter reflects the breeders’ emphasis on its importance. They demonstrated that their GM formulation is equivalent to an optimization problem of minimizing inbreeding subject to defined level of risk, denoted ρ. The solution needs to calculate risk and inbreeding for the range of acceptable ρ values for N_c_ progeny from N_p_ parents, i.e., (*N_p_C* _2_)^Nc^ / N_c_! (Akdemir and Sanchez 2016) developed a Tabu-search GA to determine the efficiency frontier between inbreeding and risk. In an updated version, (Akdemir 2018) used a GA to find the complete set of non-dominated solutions (Deb 2003, 2011) that comprise the efficiency frontier for the three criteria of Gain (G), Inbreeding (I) and Usefulness (U) values in the objective function. This allows selection of a subset of solutions for evaluation obviating the need for conducting a grid search across all possible values.

Akdemir and Sanchez (2016) demonstrated the utility of their genomic mating approach using simulations of recurrent selection beginning with two founders for a trait composed of simple additive genetic architecture. The QTL were evenly distributed across a simulated genome consisting of three diploid linkage groups. Their results indicated that the efficiency frontier can be selected to produce responses across 20 cycles that were better than PS and as good as GS and GOCS for the first five to seven cycles and better than PS, GS and GOCS thereafter (Akdemir and Sanchez 2016). They did not include WGS for comparison in their study.

Recognizing that IMGA’s are very efficient at finding global optima Yabe *et al* (2016) suggested that computational island models could be used to create efficient and effective breeding plans for plant breeders. Even though computational IMGA’s allow the software developer to change mutation and recombination rates, which are not under the control of plant breeders, structures of breeding populations based on island models could offset loss of useful genetic variability through regulation of exchange of genotypes among sub-populations. It is not unusual for plant breeders of crops that are easily self-pollinated to routinely evaluate, select and recurrently cross lines derived from one or two specific bi-parental crosses. In the vernacular of commercial soybean and maize breeders this is known as “working a population”. Yabe et al (2016) demonstrated GS on populations organized as islands of families provided greater response to selection than GS after the seventh of 20 cycles of RGS. Their founder population consisted of lines derived from *in silico* crosses of six homozygous rice lines with an elite rice variety. They isolated the six families of RIL’s for recurrent selection using GS with no or occasional exchange of selected lines among the family islands. While their results appeared to be similar to WGS, they did not compare their results with WGS. They also suggested that the trade-off between genetic gain and retention of useful genetic variance could be improved by adjusting the number and frequencies of migrants among sub-populations.

Inspired by Akdemir and Sanchez (2016) and Yabe et al (2016), we hypothesized that a breeding strategy that organized the breeding populations as island families and utilized a genomic mating MD would provide small soybean genetic improvement projects with the ability to minimize the trade-offs between maximizing genetic gain and minimizing loss of useful genetic variability. Within the IM organized populations we evaluated three migration policies among the families. For both non-isolated and island models we applied three selection methods and four mating designs. To evaluate the potential of these combinations of methods to realize genetic gains while retaining useful genetic diversity, we compare outcomes from simulated recurrent selection applied to contemporary soybean germplasm adapted to MZ II and III using a set of metrics (Ramasubramanian and Beavis 2020). The metrics include the standardized genotypic value (R_s_), the most positive genotypic value (*M_gv_*) among RIL’s selected in cycle c, the standardized genotypic variance (*SV_g_*), the average expected heterozygosity (Hs), the lost genetic potential of populations based on the number of favorable alleles that are lost.

## 2. Methods

### 2.1 Simulations

Initial sets of soybean lines were generated by simulating crosses of 20 contemporary homozygous lines representing diversity of soybean germplasm adapted to MZ’s II and III with IA3023 to generate *in silico* F1 progeny (Ramasubramanian and Beavis 2020). Individual F1’s from each of the 20 crosses were self-pollinated *in silico* for four generations to generate 100 lines per family forming populations of 2000 lines organized into 20 families with genotypic information at 4289 genetic loci (Song et al. 2017). Thus the genetic structure of the initial simulated populations is similar to that used in the experimental SoyNAM investigation (Guo et al. 2010; Takuno et al. 2012; Song et al. 2015; Song et al. 2017; Xavier A et al. 2017; Diers B et al. 2018).

As reported previously (Ramasubramanian and Beavis 2020), there were 3818 polymorphic loci in the combined population of 20 families and an average of 773 polymorphic loci within each of the families for the initial founding sets of lines. The variance among families was ∼ 34 polymorphic loci. Across the 20 families of cycle 0 (C0) lines, average expected heterozygosity was 0.09 with an estimated variance of 4.4*10^-7^ among families. The average estimated G_st_ value across the genome for the initial founding set of RILs was 0.32, as determined by the ‘diff_stats’ function in the mmod R package (Jombart 2008; Ryman and Leimar 2009; Jombart and Ahmed 2011; Ramasubramanian and Beavis 2020). Average pairwise ‘Fst’ estimated using ‘pairwise.fst’ in ‘hierfstat’ R package (Goudet 2005) among the 20 families in simulated SoyNAM data is 0.20. Pairwise ‘Fst’ is a measure of population differentiation among pairs of populations, which is estimated as the ratio of difference between the average of the expected heterozygosity of the two populations and total expected heterozygosity of the pooled populations to total expected heterozygosity of the pooled populations. Whereas the average Fst using genotypic data from SoyNAM project among 40 families is 0.09 with a maximum pairwise Fst of 0.15 and a minimum Fst of 0.007 (Ramasubramanian and Beavis 2020).

### 2.2 Combinations of Factors

We evaluated 60 combinations of factors (Table 1) that could influence responses to recurrently selected populations derived from a set of founder genomes representing the diversity of contemporary soybean germplasm adapted to MZ II and III in North America (Mikel et al. 2010; Diers et al. 2018). The treatment factors included structure of breeding populations, selection method, and mating design. The structure of the breeding populations included organizing 20 sub-populations representing the original 20 founder families, referred to as family islands (FI), and populations in which the family structures were not retained after the initial founder population was created, referred to as non-island (NI) populations.

**Table 1.** Treatment Design representing the factors that impact responses and limits of responses. The parameter values for levels of island selection specific factors were selected based on limits of responses from a larger set of simulations (2664 combinations of factors with 10 replicates per combination of factor) with migration frequency (1, 2, 3), migration size (1, 2) and migration direction (1, 2) and mating designs (HN, CR and RM) for 40, 400 and 4289QTL with 0.7 and 0.3 H.

| Factors | Levels | Values for Levels |
| --- | --- | --- |
| Population Type | 2 | Non-isolated and Island populations |
| **Island Model Selection** |  |  |
| Migration Frequency | 1 | Migration frequency of 2 corresponds to migration of lines every other cycle of selection |
| Migration Size | 1 | Migration of 2 lines per migration event (20%). |
| Migration Policy | 4 | i) Discrete Selection<br>(For DS, migration frequency, size and direction are set to '0')<br>ii) Best Island<br>iii) Random Best<br>iv) Fully connected |
| Migration Direction | 1 | i) Bi-directional |
| **Factors Common to Non-isolated and Island Populations** |  |  |
| Selection Method | 3 | i) PS ,ii) GS, iii) WGS |
| Mating Design | 4 | i) Hub Network Design<br>ii) Chain Rule<br>iii) Random mating<br>iv) Genomic mating |
| Genetic Model Parameters | 1 | 400 QTL and 0.7 H |
| Total Number of Combinations of Treatment Factors | 60 |  |
| Total Number of Simulations | 5<br>(replicates/combination of factors) | 300 |

Previously we demonstrated that development of homozygous lines for phenotypic evaluation will limit the numbers of segregating linkage blocks with effective QTL effects each cycle of selection (Ramasubramanian and Beavis 2020). Consequently, we chose to designate only 400 polymorphic marker loci as simulated QTL. The QTL were distributed uniformly among the SNP loci and each contributed equal additive effects of ±0.5 units to the total genotypic value of a line. Thus, cycle C0 lines derived from the founders had an average genotypic value of zero and the potential to create genotypic values ranging from -200 to +200. Phenotypic values were simulated by adding non-genetic variance sampled from an N (0, σ) distribution to the simulated genotypic values, where σ was determined by the heritability. Herein we report only simulated broad sense heritability values on an entry mean basis of 0.7. The non-genetic variance was held constant across subsequent cycles of selection. Thus, heritability is expected to decline with every cycle of selection due to the loss of additive genetic variance.

Phenotypic selection (PS), genomic selection (GS) and weighted genomic selection (WGS) were applied recurrently to both population structures. Recurrent selection applied to the non-island populations consisted of ranking all lines in a given cycle (Figure 1) according to the selection metric and retaining 10% for crossing to create the next cycle of lines. In terms of standardized selection differential, this corresponds to a selection intensity, *ι* = 1.75. For selection of lines organized into FI’s, 10% of the lines are selected within FI’s (Figure 2). Subsequently, 20% of lines might be migrants from other FI’s depending on migration rules (Table 1). Metrics used for selection include simulated phenotypic values for PS, genome estimated breeding values (GEBVs) for GS and weighted genome estimated breeding values for WGS. We used the weighting function used by Jannink (2010) for estimating weighted genome estimated breeding values. The weighting functions are provided in Supplementary Table 1. Previous results indicated that among GS methods, Ridge Regression (RR) provided the best compromise between short term and long term responses (Ramasubramanian and Beavis 2020), thus we only used RR to train GP models for GS. RR was implemented with a method that employs Expectation Maximization to obtain Restricted Maximum Likelihood Estimates of marker effects (Xavier 2019).

**Figure 1.**
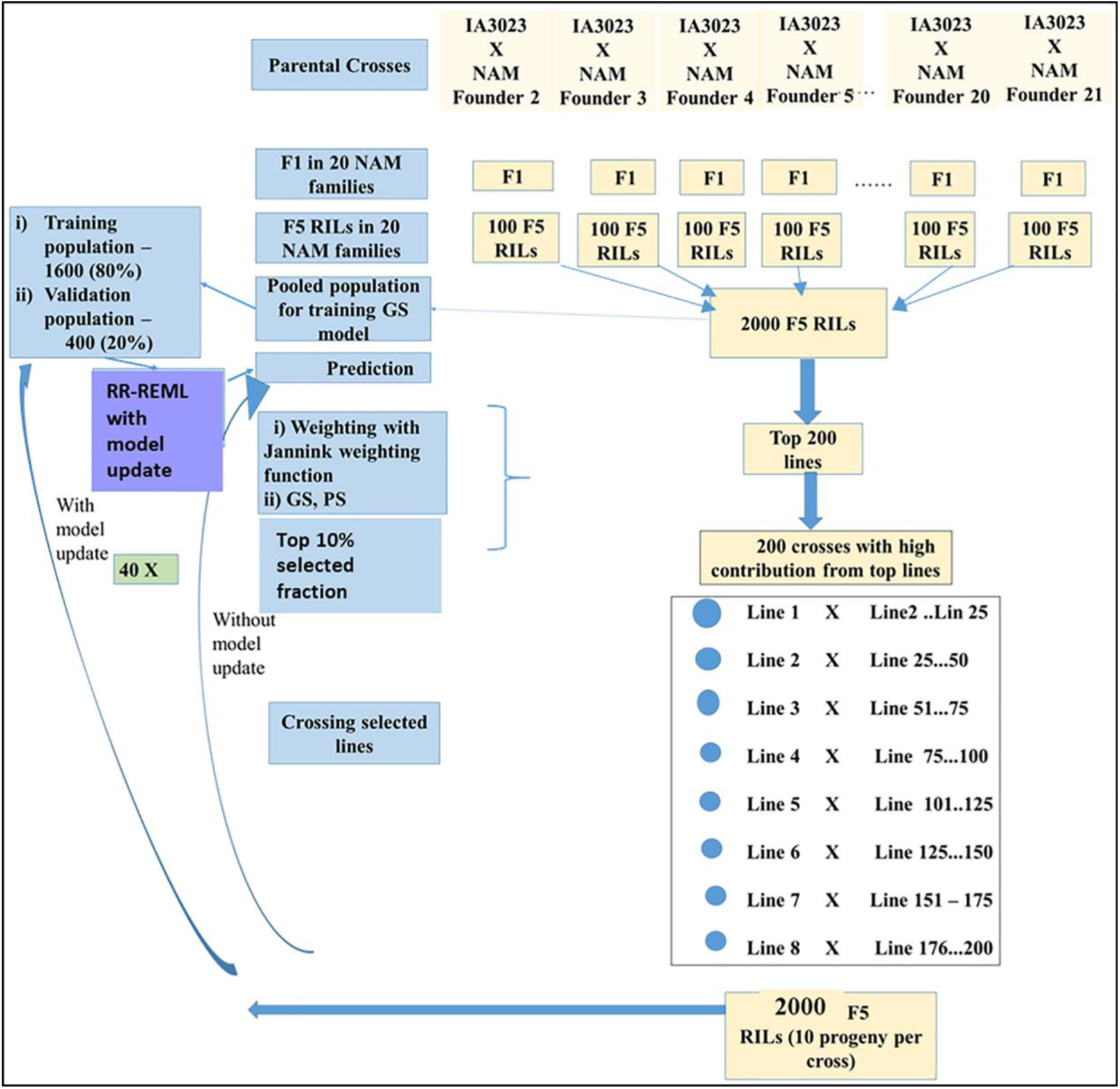
Schematic representing simulated recurrent selection in Non-Isolated (NI) populations comprised of twenty families from Soybean NAM founders. The schematic depicts the *in silico* steps used to generate the base population of 2000 F_5_ RILs derived from 20 NAM founder lines crossed to IA3023. The depiction includes the model training step and the recurrent steps of prediction, sorting, truncation selection, crossing, and generation of 2000 F_5_ RILs for each cycle as well as the decision steps to check if the training set should be updated and if the recurrent process should be continued for another cycle.

**Figure 2.**
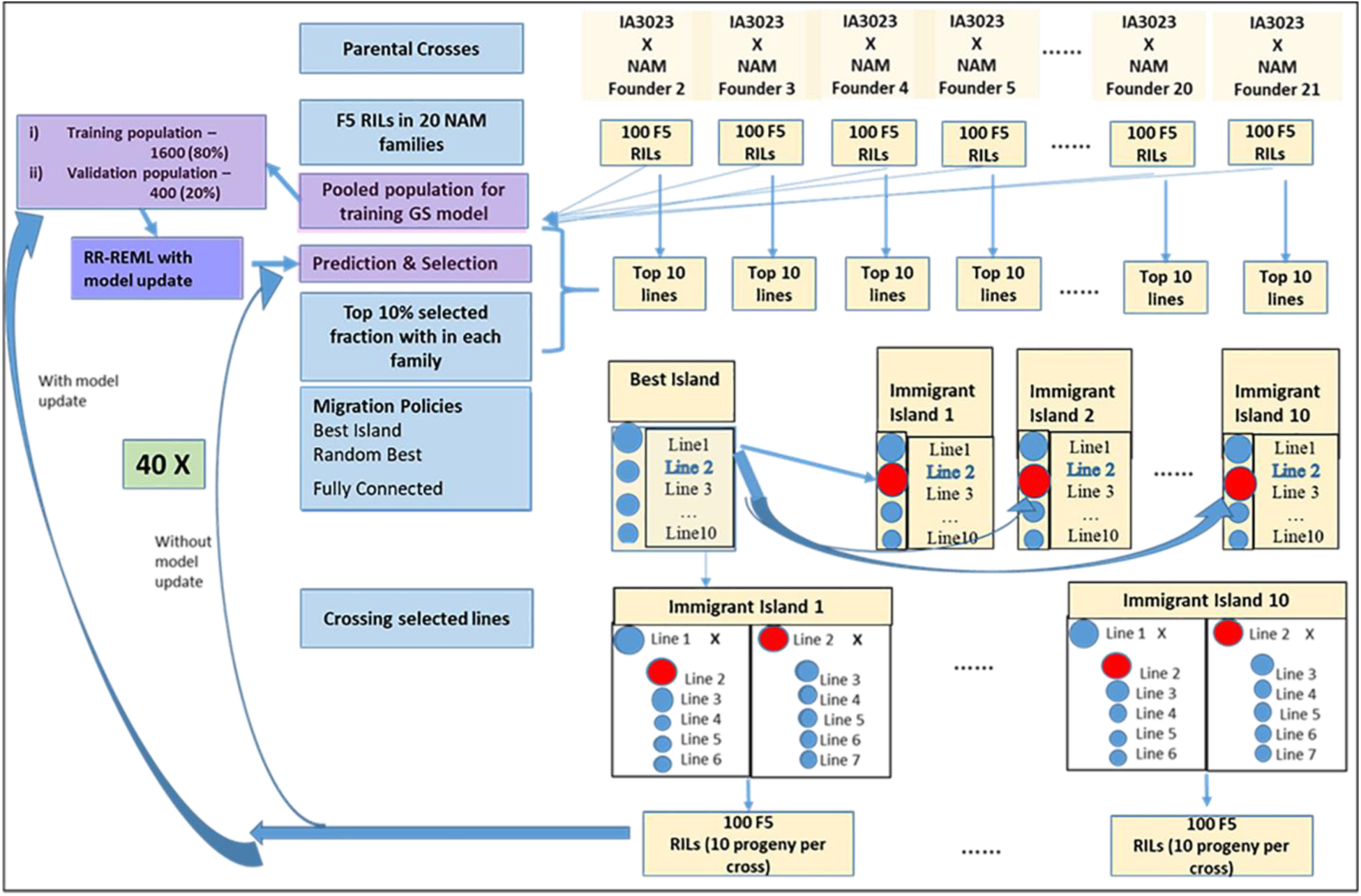
Schematic representing simulated recurrent selection of Family Island (FI) populations where each of the twenty families from the Soybean NAM founders is considered an island population. The schematic depicts the *in silico* steps used to generate the base population of 2000 F_5_ RILs derived from 20 NAM founder lines crossed to IA3023. 100 F5 RILs generated from each of the crosses form a distinct island. The depiction includes the model training step and the recurrent steps of prediction, sorting, truncation selection within islands, migration, crossing, and generation of 200 F_5_ RILs per island for each cycle as well as the decision steps to check if the training set should be updated and if the recurrent process should be continued for another cycle. The blue shaded circles represent lines that are descendants of the founder populations in the islands and red shaded circles represent lines that are replaced by immigrants from the island with the largest genotypic value for the ‘Best Island’ policy.

For both GS and WGS the training models were updated every cycle of selection with data sets from all prior cycles. Since average within family prediction accuracies are lesser than prediction accuracies from a combined TS comprising of RILs from all the families (Ramasubramanian and Beavis 2020), we used a combined TS comprising of RILs from all the families. Training sets for each cycle were obtained by randomly sampling 1600 lines from the set of 2000 lines for each cycle. The most accurate predictions and maximum genetic responses were obtained with training data that is cumulatively added every cycle. For purposes of this manuscript, model updating refers to retraining the model with data from the current cycle as well as all prior cycles that were cumulatively added.

Subsequent to selection, four mating designs were applied to create the next cycle of lines (Table 1). To simulate theoretical truncation selection, selected lines were randomly mated (RM). The chain rule (CR), a.k.a., a single round-robin mating design (Yabe et al. 2016), is an alternative to RM that assures all selected lines contribute to the subsequent cycle of evaluation and selection. In contrast to the attempt to assure equal representation of selected lines through RM and CR, most soybean breeders use a mating design that assures most progeny will be derived from crosses of a few lines that exhibit the most desirable performance (Guo et al. 2013; Guo et al. 2014). Because the metaphor of hubs with spokes represents the preference for crossing most selected lines to a few “hub” lines, we refer to this mating design as a hub network (HN) and is the mating design used in our previous investigation (Ramasubramanian and Beavis 2020). The fourth mating design, genomic mating (GM), uses mathematical objective functions to assure that defined breeding objectives are used to identify pairs of crosses from among the selected lines. Genomic Mating was implemented with the ‘Genomic Mating’ R package (Akdemir et al. 2018). As originally described GM combines selection and mating in a single step, but we decomposed the steps to provide comparable outcomes from all other combinations of selection methods, mating designs and organized populations.

#### 2.2.1 Genomic mating in non-isolated families

In a selected set of 200 lines there are 200C2 (19900) combinations of parental pairs. To solve the objective function w.r.t an initial population of parental pairs, 250 initial populations of 200 combinations of parental pairs are sampled from 19900 combinations (19900C200) for the GA algorithm to solve.

#### 2.2.2 Genomic mating in populations organized as family islands

In island selection, ten lines are selected from each of the 20 family islands. Within each island 45 (10C2) combinations of parental pairs are possible (Supplementary Figure1). To solve the objective function w.r.t an initial population of parental pairs, 250 initial populations of 10 combinations of parental pairs are sampled with replacement to keep the population size equal to the NI populations for the GA algorithm. For each of the 20 families, the GA algorithm is applied to the initial subset of 250 out of all possible combinations (45C10). The other parameters for the GA algorithm are the same for both NI and FI populations. The GA algorithm selects non-dominated elite solutions (Deb 2003, 2011) and mates of non-dominated elite solutions for 50 iterations with a mutation probability of 0.8 (Supplementary Figure1). Examples of pseudocode are provided in (Akdemir and Sanchez 2016) and the Genomic Mating R package manual (2018). It is important to note that the parameters values in the GA algorithm can be optimized and the set of solutions in the pareto-front can be explored for better solutions by other methods such as NSGA-II, NSGA-III, SPEA-1, SPEA-2 and other recent improved versions of GA for better convergence rate and quality of solutions, determined by the proximity to global optimum (Deb 2011; Seada and Deb 2018) (Supplementary Figure1).

### 2.3 Migration Rules among family islands

In addition to applying selection methods and mating designs to both population structures, there are many possible rules that affect migration among islands. A preliminary investigation of migration rules that was implemented included: 1) Frequency of migration - never, once every two cycles and every cycle of recurrent selection. 2) The proportion (10% and 20%) of immigrants that will be included in crosses responsible for creating the next cycle of lines. 3) Migration can be either in one direction or it can be reciprocal among family islands. Based on the preliminary investigation, we decided to set migration rules bi-directional migration between both immigrant and emigrant islands of two lines once every other cycle of selection.

#### 2.3.1 Migration Policies among family islands

included three levels included “Best Island” (BI), “Random Best” (RB), and “Fully Connected” (FC). Migration policy (MP) refers to the nature of island topology specifying connections between emigrant and immigrant islands. For the BI policy, emigrant lines are selected from the island with most desirable genotypic value and the selected lines can emigrate to no more than 10 islands. Given a bi-directional migration rule, the emigrant island also receives two immigrants from the islands that received the emigrants. For a RB policy, an emigrant island is selected randomly from a set of islands with high genotypic values, while the migration pattern itself is similar to BI policy. For the FC policy, every island is connected to every other island and lines migrate from emigrant islands with high values to randomly selected immigrant islands (Supplementary Figure2).

Note that migration factors are irrelevant for populations that did not maintain the structure of FIs and they are irrelevant for FI’s that do not experience migration. Thus the treatment design is not a complete factorial, rather the complete set is comprised of responses for 60 combinations of factors.

### 2.4 Modeled response to recurrent selection

The averaged genotypic values *y*, for each cycle, c, of recurrent selection were modeled with a linear first order recurrence equation:

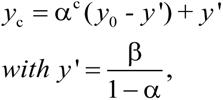

Where *y*_0_ specifies the average genotypic value of the lines derived from the founders, α^c^ represents the difference in response between the current cycle and the previous cycle, *y*’ represents the asymptotic limit to selection and c consists of a sequence of integers from 0 to 39 representing each cycle of recurrent selection (Goldberg 1958; Ramasubramanian and Beavis 2020). The parameters *y*_o,_ α, and β were estimated with a non-linear mixed effects method implemented in ‘nlme’ functions in the ‘nlme’ and ‘nlshelper’ packages (Pinheiro and Bates 2000; Baty et al. 2015; Pinheiro et al. 2019).

For asymptotic limits of responses, the number of cycles required to reach half of the limits before there is no longer response to selection is referred to as the half-life of the recurrent selection process (Robertson 1960; Dempfle 1974; Kang 1979; Cockerham & Burrows 1980; Kang and Namkoong 1980; Kang 1987; Kang and Nienstaedt 1987). From the first order recurrence equation, the half-life is estimated as ln (0.5)/ln (α), when y0 is ‘0’ and the asymptotic limit is estimated as y’ (Ramasubramanian and Beavis 2020).

### 2.5 Analyses of variance (ANOVA) of modeled response to recurrent selection

ANOVA is used to evaluate the impact of factors and their interactions on the modeled responses to global and island recurrent selection. The analyses of variance used single level nlme models with modeled (eqn 4) responses grouped by combinations of treatment factors. We analyzed the variance among modeled responses using AIC, BIC and Likelihood metrics that were grouped based on combinations of treatment variables consisting of population type, selection method, mating design, and migration policy for a constant level of migration frequency, migration size and migration direction for one genetic model consisting of 400 simulated QTL responsible for 0.7 H with equal additive effects (Table 1). For a discussion of ANOVA using non-linear mixed effects models refer (Pinheiro et al 2000; Zuur 2009; Baty, et al. 2015; Pinheiro et al, 2019; Oddi et al. 2019; Ramasubramanian and Beavis 2020).

In the first phase of model fitting, we fit a random intercept model for estimating both alpha and beta in the recurrence equation using the ‘nlme’ R package. Estimates of modeled parameters from nlsList models were retained as starting values for fixed effects. Multiple ANOVA of ‘nlme’ objects representing the models were used to identify combinations of factors with significant effects on the non-linear response. The model with the lowest AIC score was selected as the best model. The best random intercept model in the first phase of model fitting process M15 and models with combinations of three factors (M11-M14) showed auto-correlation of residuals. Since auto-correlation violates the independence assumption, the correlation among residuals was modeled using AR-1 correlation structure. Since the genotypic values across cycles in recurrent selection are correlated, fitting AR-1 correlation doesn’t remove the correlation unless cycles are used as co-variates. However, using cycles as a co-variate makes the model fitting very time consuming and often has larger AIC scores than models without cycles as co-variates. The Model M15 with AR-1 correlation structure was further refined by modeling variance components using ‘varIdent’ structure in ‘nlme’. The process of fitting, selecting and refining mixed effects models is similar to our previous study (Ramasubramanian and Beavis 2020) and is described in the vignettes in R package ‘SoyNAMSelectionMethods’.

### 2.6 Evaluations of Responses to Recurrent Selection

were conducted on both modeled and genotypic values using a set of metrics described in (Ramasubramanian and Beavis 2020) and defined below. The estimated population half-life and asymptotic limits used the estimated parameters, α and β of the first order recurrence model. The average genotypic values were used to estimate the standardized genotypic value (Rs) and maximal genotypic value (Mgv).

Maximum possible genotypic potential of the founders provided a reference for number of favorable alleles retained in the population. The loss of genotypic potential is characterized by reduction in the standardized variance of genotypic values (Sgv) and estimated heterozygosity (Hs). In addition, efficiency of conversion of loss in genotypic variance into genetic gain (Rs_var) provides a way to assess gain in genotypic value and loss of genetic variance simultaneously. In island model selection, the different impacts of selection strategies on the genotypic variance at island or global levels are assessed using intra-island Sgv, inter-island and global variance of genotypic values. A schematic diagram of the factors and evaluation metrics used to characterize the responses to recurrent selection is provided in Figure 3.

**Figure 3.**
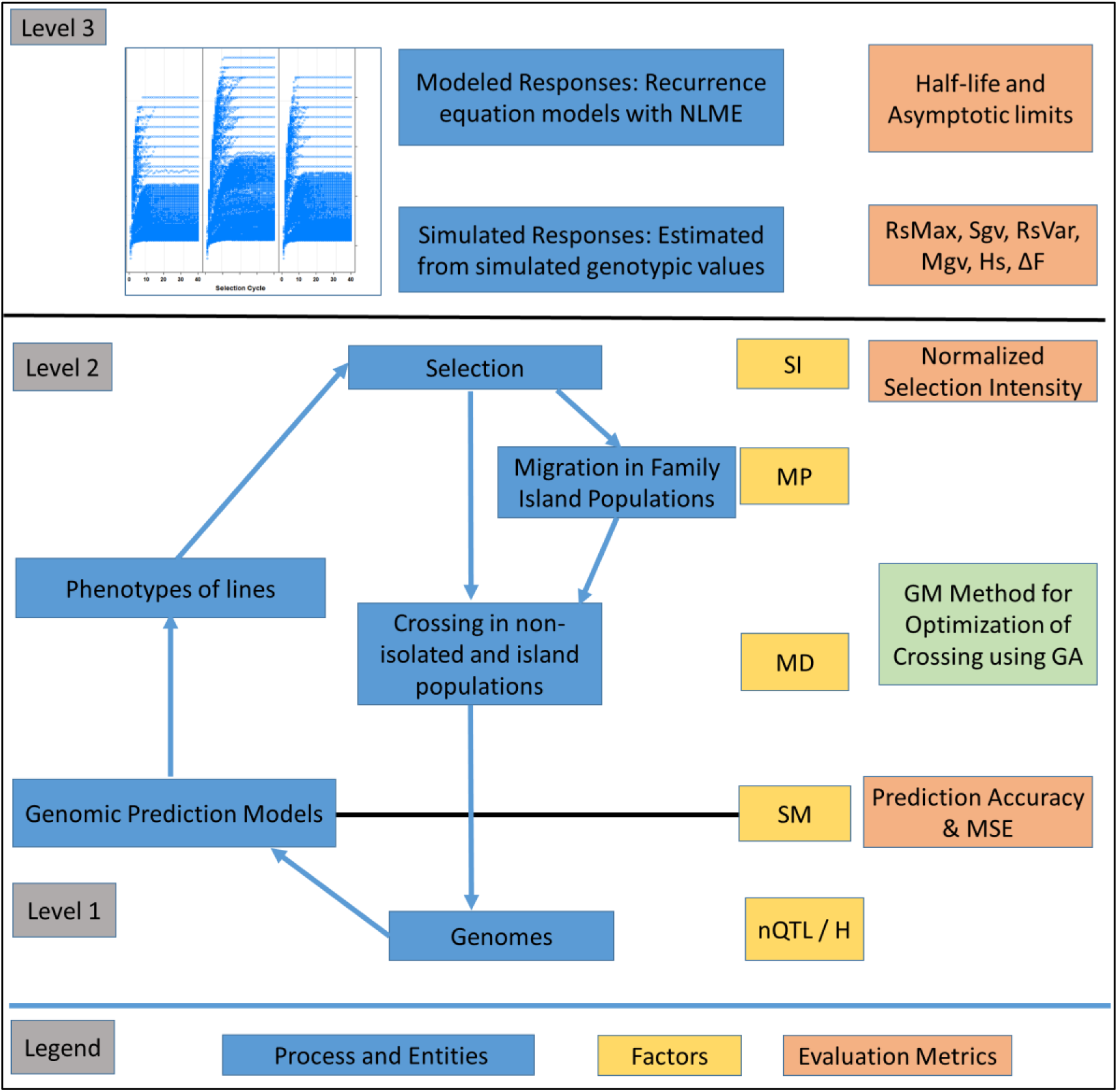
Overview of the Recurrent Selection Process: Representation of entities such as genomes, associated RILs and processes such as the estimation of marker effects, selection, migration and crossing. Levels correspond to layers of information with level 1 comprised of genomic information, level 2 comprised of phenotypes of lines within and across families and level 3 comprised of higher level information including responses across cycles of selection. The factors include nQTL and H at the genome level, SM (selection method) including PS, GS and WGS. The factors at level 2 includes SI (top 10% selected fraction), MD (Mating design, which includes Hub Network, Chain Rule, Random Mating, and Genomic mating levels) and MP (Migration Policy, which includes “Best Island”, “Random Best” and “Fully Connected” policies). Among the BD levels, GM method involves application of evolutionary multi-objective optimization to minimize inbreeding and maximizing gain. Level 3 is characterized using evaluation metrics such as half-life and asymptotic limits derived from recurrence equation models and metrics such as standardized responses (Rs), Standardized genetic variance (Sgv), Maximal genotypic values (Mgv) and efficiency of converting loss in genetic variance into gain (RsVar) derived from simulated outcomes. Other metrics include prediction accuracy and MSE for GP models (RR-REML) and expected heterozygosity (Hs).

#### 2.6.1 Evaluation Metrics

The standardized genotypic value, R_s_ (Meuwissen et al. 2001; Liu et al. 2015; Ramasubramanian and Beavis 2020), was estimated every cycle of selection as the proportion of maximum genotypic potential (200 units) relative to the average genotypic value of 2000 lines in C0 (1). Values range from 0-1 with the value of 1 corresponding to the maximum possible genotypic value with the genetic model and 0 corresponding to the average genotypic value of C0.

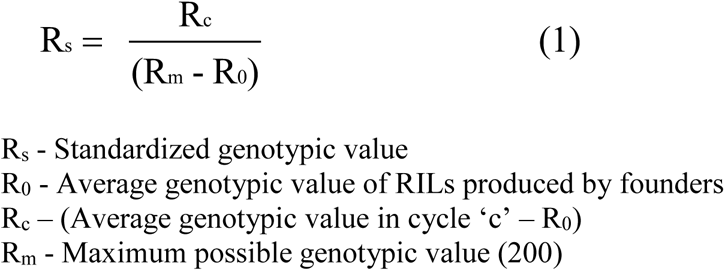

Since we previously evaluated genetic improvement of soybean using PS and the HN mating design in NI populations, we used PS with a selection intensity of 1.75 for NI population and HN mating design (designated as NI-PS-HN) as a reference for comparing other selection and mating designs. A standardized relative genotypic response, ΔRs_c_ is calculated in equation (2) as the percentage of the difference in standardized genotypic values, Rs_c_, in each cycle c.

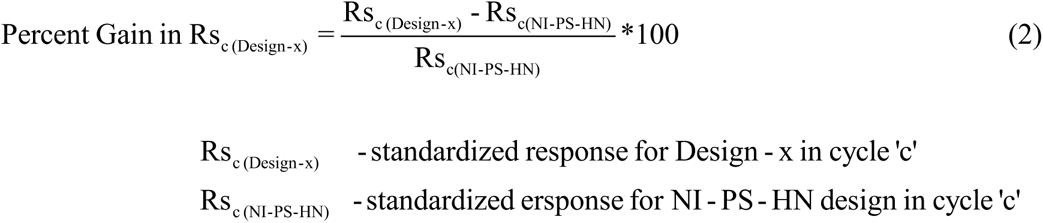

The standardized genotypic variance (*Sgv*) defined as the estimated genotypic variance divided by the estimated genotypic variance of the initial sample of lines from C0 was used to evaluate the changes in estimated genotypic variance across cycles of recurrent selection. Note that values for the *Sgv* range from zero to one.

Efficiency of genetic improvement is a metric used to estimate the proportion of genetic improvement that was obtained through loss of genetic diversity from recurrent selection (Gorjanc et al. 2018). Efficiency is estimated as the slope in linear regression in linear regions of response curves. However, responses to recurrent selection in the absence of mutation are inherently non-linear (Robertson 1960; Hill and Robertson 1968; Bulmer 1976; Ramasubramanian and Beavis 2020). For purposes of evaluating the relative contribution of lost genetic variance to genetic response in both linear and non-linear segments of the response curve, we introduce the standardized genotypic variance of the response, Rs_Var, calculated with equation (3).

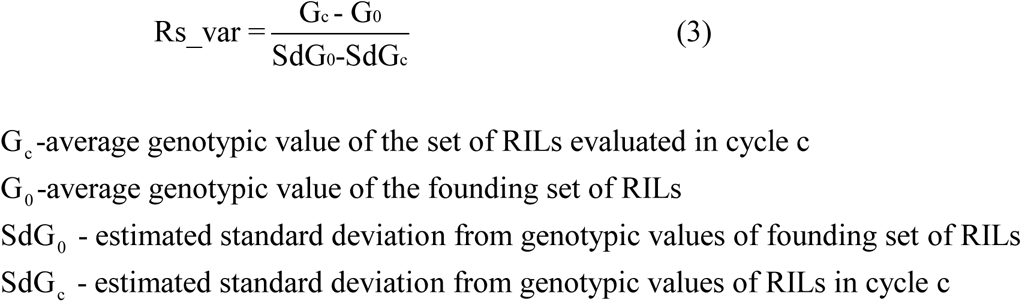

The numerator term represents difference in average genotypic values of a population in cycle ‘c’ from cycle ‘0’ normalized to standard deviation of genotypic values in cycle ‘0’. The denominator represents difference of standard deviation of genotypic values between cycles ‘0’ and cycle ’c’ normalized to the standard deviation of genotypic values in cycle 0 (Ramasubramanian and Beavis 2020). For the NI populations, Rs_Var was estimated by calculating the variance of simulated genotypic values. Standardizing the estimated genotypic variance with respect to the maximum genotypic values in the initial population, results in values that range from 0-1. For the FI populations, the genotypic variances can be split into within and between island genotypic variance. The three measures we used to estimate the global diversity of populations, inter-island diversity and within island diversity are provided in the documentation of the R package.

## 3. Analyses and Data Availability

Simulated data and software codes are available as part of the R package ‘SoyNAMSelectionMethods’ (Supplementary File1). Documentations for downloading and using the package are available at http://gfspopgen.agron.iastate.edu/SoyNAMSelectionMethods_v2_2020.html. The SoyNAM founder genotypic and phenotypic data are available in SoyBase (Grant et al., 2010).

## 4. Results

### 4.1 Rates and limits of responses to recurrent selection

Factors common to NI populations and FI populations such as mating design and selection method as well as factors specific to discrete and island model selection had significant effect on estimated population half-lives and asymptotic limits. Half-lives for selection methods on NI populations ranged from 3.83 to 16.10 cycles with a mean of 9.62 cycles and asymptotic limits ranged from 71.64 to 160.76 with a mean of 115.97 (58% of the maximum possible potential in the founders). Compared to NI populations, half-lives for discrete selection (DS) methods were very low ranging from 1.97 to 2.89 cycles with a mean of 2.43 cycles and asymptotic limits ranged from 28.42 to 38.30 and a mean of 33.12 (16.5% of the maximum possible potential in the founders) (Supplementary File2; Supplementary Figure3). Estimated half-lives for island model selection methods were greater, on the average than NI methods ranging from 4.24 – 32.04 cycles with a mean 13.45 cycles and asymptotic limits ranged from 47.54 to 198.82 with a mean of 116.8 (58.5 % of the maximum possible potential in the founders) (Supplementary File2; Supplementary Figure3).

### 4.2 ANOVA of modeled genotypic values

There is strong evidence from the analyses of variance (Supplementary File3) that the modeled genotypic values across cycles of selection depend on interactions among selection method, mating design and migration policy. The most parsimonious model included all combinations of factors indicating interactions among all factors have statistically significant influences on recurrent responses to selection and requires unique estimates of α, and β in (3) for each of the combinations of factors (M15 in Supplementary File3). For all combinations of factors, we report only migration involving bi-directional migration of two migrants every other cycle. Among the factors that affect only FI populations, migration frequency had significant effects on rate and the asymptotic limits for response to selection, whereas migration direction and size had relatively small effects on rates and no significant effect on the asymptotic limits for response to selection. Rates and genotypic values at the limits of response for a given selection method and mating design also depend on genetic architecture and heritability (data available on request). Rather than belabor the specific outcomes from all possible combinations of factors that affected the modeled responses, the remainder of the reported results are restricted to results from simulations with 400 QTL responsible for 70% of phenotypic variability.

### 4.3 Responses to recurrent selection of Non-Isolated lines

There were 12 combinations of selection methods and mating designs that were applied to lines of NI populations. The greatest genotypic values (Rs) were attained with WGS (Figure 4 and Supplementary Figure 4). Genomic selection using RRBLUP values resulted in greater responses than PS in early cycles while WGS produced greater responses than PS in later cycles (Figure 4; Supplementary Figure4). Weighted genomic selection followed by the CR mating design resulted in the greatest realization of genetic potential before reaching a limit. Genomic selection using RRBLUP values followed by a hub network (HN) mating design resulted in the greatest rates of response in the first ten cycles and if followed by RM, provided the greatest responses in the first 20 cycles. When the GM design is applied to selected lines to obtain specified crosses according to optimization criteria, the responses in the first 15 cycles were larger than obtained with RM, whereas responses after the 20^th^ cycle were less than responses for other mating designs (Figure 4 and Supplementary Figure 4).

**Figure 4.**
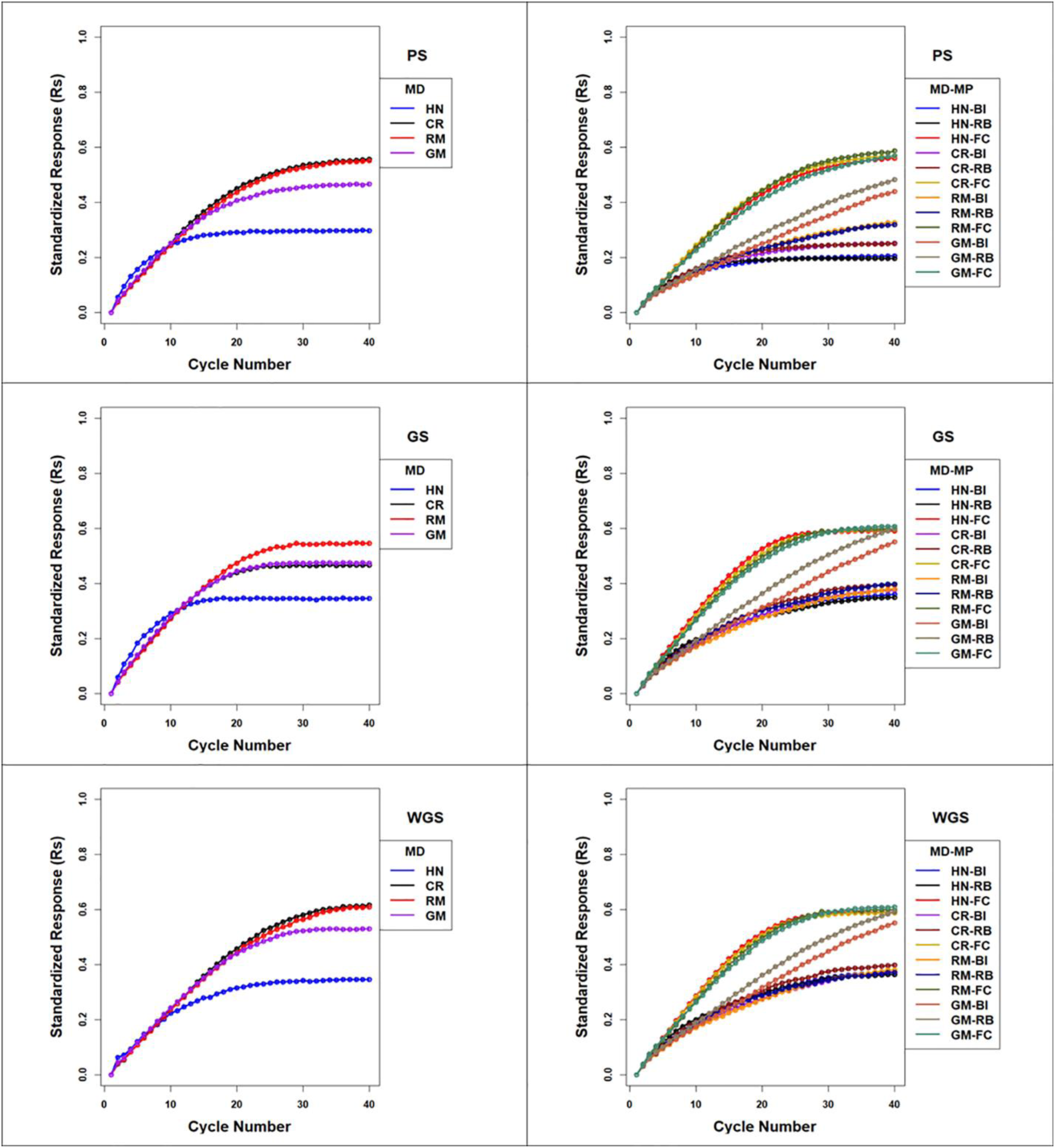
Standardized Genotypic Responses (Rs) across 40 cycles of recurrent selection on non-isolated (left panels) and family island (right panels) populations, using PS (top panels), GS (middle panels) and WGS (bottom panels) for the four mating designs: Hub-Network (HN), Chain Rule (CR), Random Mating (RM), and Genomic Mating (GM). Standardized genotypic responses are represented from a simulated genetic architecture consisting of 400 additive QTL uniformly distributed throughout the genome and responsible for 70% of phenotypic variability. Ten percent of lines are selected to be used in crosses in HN, CR, RM and GM designs. Migration policies include bi-directional migrations of two migrants every other cycle involving the Best Island (BI), Random Best (RB), and Fully Connected (FC) migration policies.

**Figure 5.**
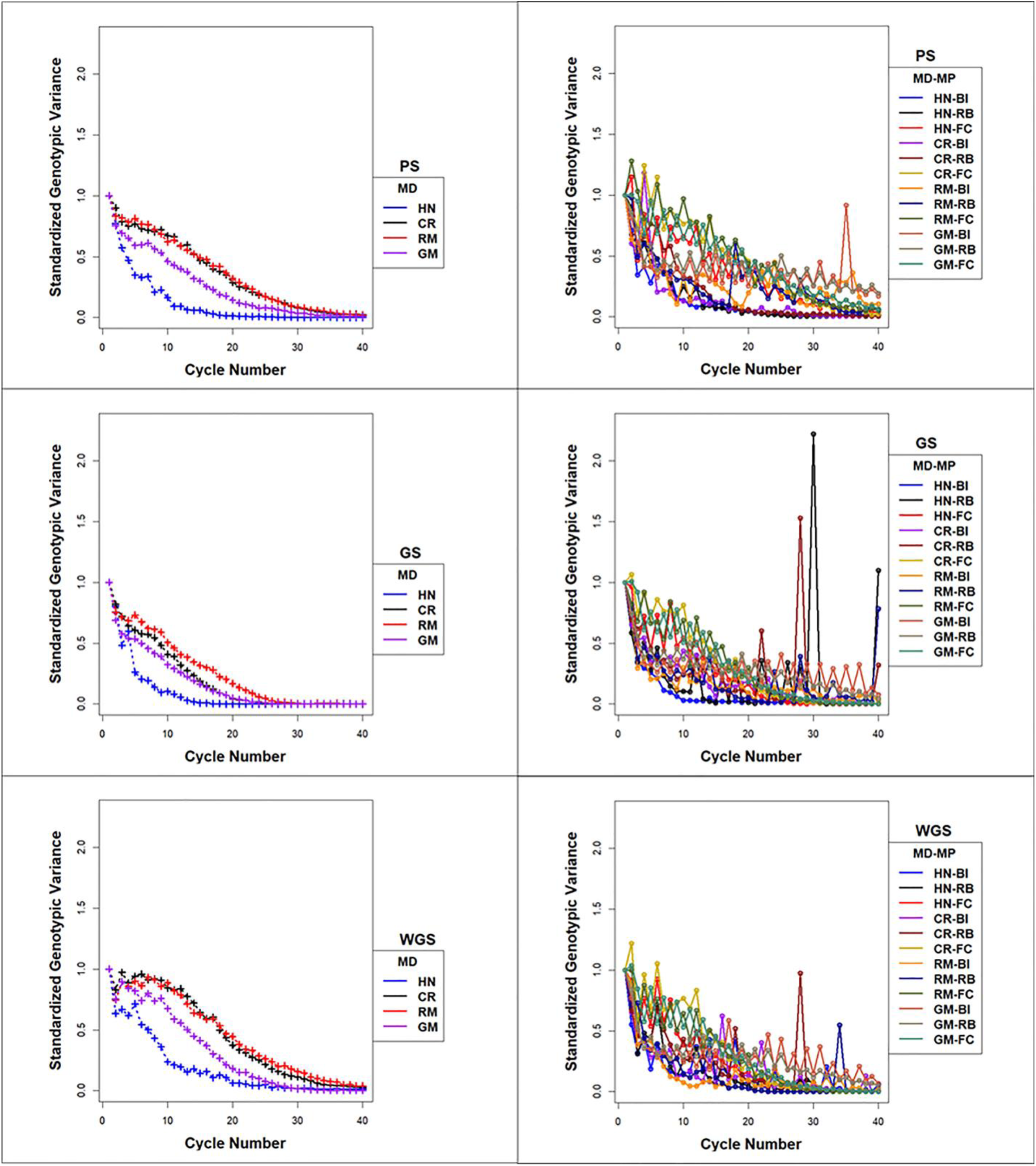
Standardized Genotypic Variance across 40 cycles of recurrent selection on non-isolated (left panels) and family island (right panels) populations, using PS (top panels), GS (middle panels) and WGS (bottom panels) for the four mating designs: Hub-Network (HN), Chain Rule (CR), Random Mating (RM), and Genomic Mating (GM). Ten percent of lines are selected for crossing. The genetic architecture in the initial simulated founder lines consisted of 400 additive QTL uniformly distributed throughout the genome and expressed broad sense heritability on an entry mean basis of 0.7. Genetic variance is standardized to the average genotypic variance in founder populations in cycle ‘0’. Average island genetic variance refers to genetic variance within families averaged across 20 families. Migration policies in the island models consisted of bidirectional exchange of two immigrants and emigrants every other cycle of selection. Migration policies include BI- “Best Island”, RB- “Random Best”, and FC- “Fully Connected”

The responses measured as maximum genotypic values (Mgvs) produced response patterns similar to Rs. Use of WGS followed by the chain rule (CR) mating design resulted in an average Mgv of 125 (62.5% of the maximum potential in the founders) followed by PS and GS using RRBLUP values in the 40^th^ cycle. Genomic selection followed by the HN mating design (NI-GS-HN) realized greater Mgvs relative to other combinations of factors only in the early cycles (Supplementary Figure 5).

The rate at which maximum genotypic potential decreased across cycles of selection was reflected in the estimated number of lost favorable alleles. Among the selection methods, GS using RRBLUP values lost genetic potential faster than PS and WGS. Among the mating designs, HN resulted in the fastest loss of genetic potential while RM lost genetic potential slower than any of the other mating designs. Genomic mating lost genetic potential at a rate that was intermediate between RM and HN mating designs. The CR design lost favorable alleles at rates that were similar to GM after GS, whereas after applying CR after PS and WGS, the loss of alleles was similar to RM (Figure 4).

The rates at which favorable alleles were lost exhibited similar patterns as the changes in standardized genotypic variance (Sgv) and expected heterozygosity (Hs) (Figure 4 and Supplementary Figure 6). The application of RM and CR mating designs after selection helped maintain genotypic variance and heterozygosity for use in later cycles of recurrent selection. The HN mating design resulted in the fastest loss of Sgv and Hs (heterozygosity) while the GM design demonstrated losses of Sgv and heterozygosity that were intermediate between HN and RM/CR designs.

Rates of inbreeding are larger for GS compared to PS and WGS in the first 10-15 cycles. The RM and CR mating designs demonstrated the slowest rates of inbreeding, whereas inbreeding with the GM and HN mating had high rates of inbreeding before responses to selection became limited (Supplementary Figure 7 & 8). The estimates of genotypic responses, standardized to genotypic variance (Rs_Var), were the greatest in the first 20-30 cycles with CR, RM and GM mating designs while the HN mating design lost the greatest amount of phenotypic variance after GS, PS and WGS (Supplementary Figure 9 & 10).

### 4.4 Responses to recurrent selection of lines organized as Family Islands

The genotypic values when the population reached a limit using Discrete Selection (DS), where there is no exchange of lines between islands, were as much as 67% less than the values when limits were reached in the non-isolated counterpart populations (Supplementary Figure 11). Among the DS methods, GS and WGS with GM design (designated DS-GS-GM and DS-WGS-GM) provided the greatest genotypic values at the response limits. Between 10-15% of the maximum potential in the founder populations were realized within the first 10-15 cycles with DS (Supplementary Figure11). Mgvs followed a pattern similar to Rs, and Sgvs mirrored the response pattern in DS (Supplementary Figure 11).

In contrast to isolated family islands (FIs) with DS, genotypic values at the limits to selection responses were larger using BI, RB and FC migration policies among islands. Among the selection methods applied to the FI populations, GS and WGS realized the greatest genetic potential before reaching limits of responses. The impacts of mating designs on the responses to selection applied to FI populations are distinct from mating designs in NI populations. In the NI populations RM and CR mating designs provided the greatest genotypic values before response to selection became limited, whereas in the FI populations GM provided the greatest genotypic values when coupled with BI and RB migration policies. The fully connected (FC) migration policy with the largest migration rates, produced responses that were similar among the HN, CR, RM and GM designs.

As noted above, the best responses to selection in the first 10 to 20 cycles of NI populations were obtained using GS followed with a HN or GM mating design (respectively designated NI-GS-HN and NI-GS-GM in Figure 4). The greatest short-term responses to selection in FI populations were obtained using either GS or WGS followed by the HN mating design coupled to a FC migration policy (IM-GS-HN-FC and IM-WGS-HN-FC in Figure 4 and Supplementary Figure 4). Only slightly slower rates in the first 10 to 20 cycles were obtained using GS and WGS followed by the GM design coupled to a FC migration policy.

Given a FC migration policy, the largest standardized genotypic responses at the limits to response (0.59 – 0.61) were obtained using GS or WGS with HN, CR, RM and GM designs Given a RB migration policy, GS and WGS followed by GM design produced the greatest realization of genetic potential before the 40^th^ cycle (0.59 -0.6) compared to (0.3-0.4) with HN, CR and RM designs (Figure 4 & Supplementary Figure 4). The BI policy showed a pattern similar to that of RB, but at a slower rate of response (Figure 4 & Supplementary Figure 4). Mgvs followed a pattern similar to Rs for most of the island selection methods. In contrast to selection in NI populations where PS and WGS resulted in the greatest Mgvs in 20-40 cycles, GS in FI populations resulted in larger Mgvs (124.6) than island PS (119.9) by the 40^th^ cycle.

Rates of decrease in maximum available potential is influenced by factors such as selection intensity, SM and MD. Relative to NI populations, island selection retains allelic diversity in the combined population as selection depletes variance only within islands and not across islands (Figure 6). Such loss in maximum potential is not always reflected in rates of responses. Relaxed selection intensity will result in retention of genetic variance with no significant increase in response as it is observed with BI and RB migration policies when combined with RM designs for PS, GS and WGS.

**Figure 6.**
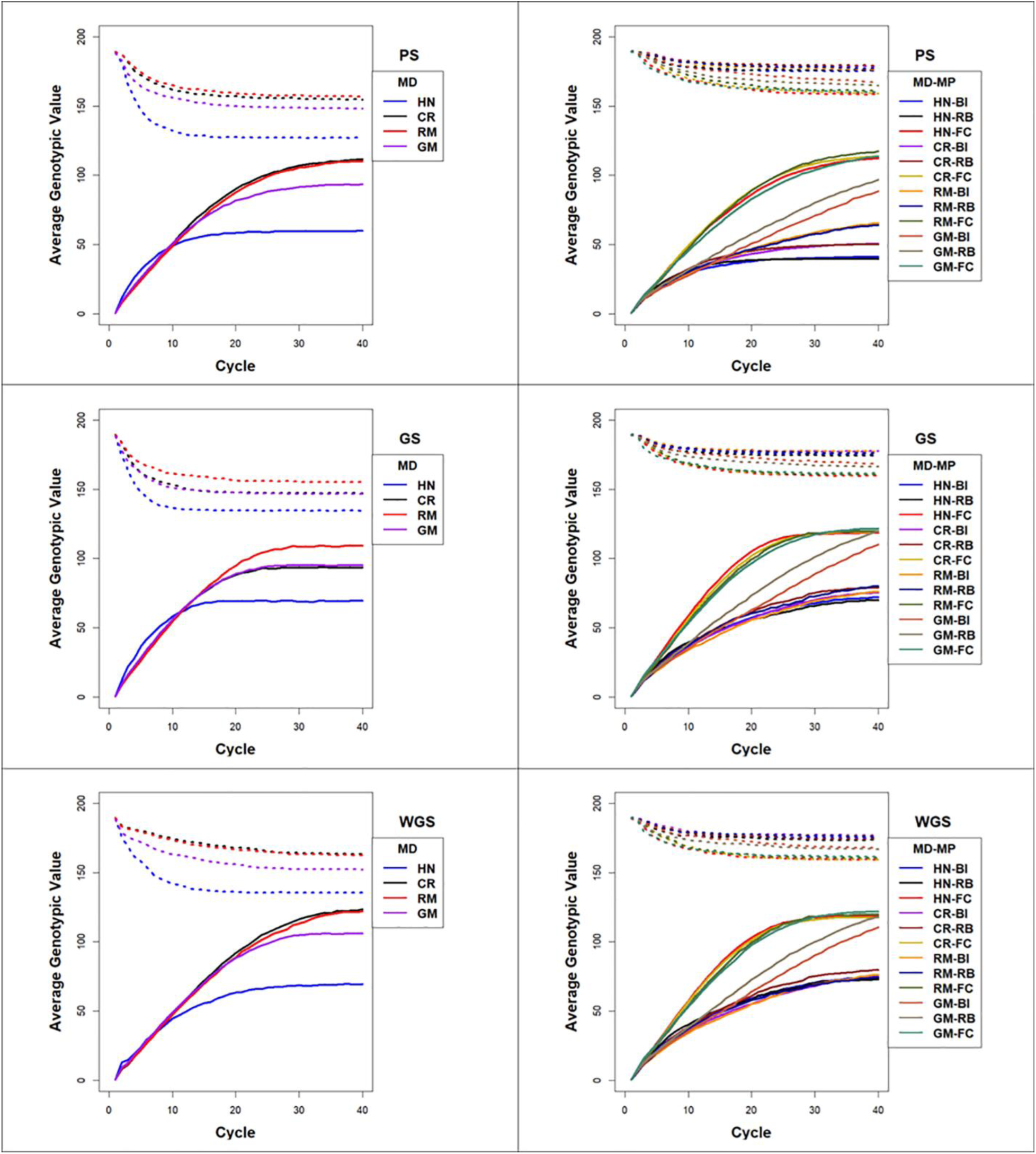
Lost Genotypic Potential and Average Genotypic Values across 40 cycles of recurrent selection on non-isolated (left panels) and island (right panels) populations, using Phenotypic Selection (PS-top row of panels), Genomic Selection (GS-middle row of panels) and Weighted Genomic Selection (WGS-bottom row of panels) and four mating designs: Hub-Network (HN), Chain Rule (CR), Random Mating (RM), and Genomic Mating (GM). Ten percent of lines are selected for crosses using HN, CR, RM and GM mating designs. The genetic architecture in the initial simulated founder lines consisted of 400 additive QTL uniformly distributed throughout the genome and expressed broad sense heritability on an entry mean basis of 0.7. The dotted lines represent maximum genetic potential estimated from favorable alleles that are lost from the population and solid lines represent increase in average genotypic value of populations due to recurrent selection. Migration policies in the island models consisted of bidirectional exchange of two immigrants and emigrants every other cycle of selection. Migration policies include BI- “Best Island”, RB- “Random Best”, and FC- “Fully Connected”

IM-GM-FC design showed the least rate of decrease of Hs values for PS, GS and WGS reflecting a greater potential retained in the population followed by IM-GM-RB and IM-GM-BI designs. IM-HN-BI and IM-HN-RB designs showed the most rapid decrease in Hs across 40 cycles of selection, whereas CR and RM designs with RB and BI migration policies showed intermediate rates of decrease of Hs. There is an oscillatory pattern in the decrease of Hs, where Hs increased with every migration event in early cycles. In late cycles, the magnitude of increase in Hs due to migration event decreased and the oscillatory pattern dampened to a continuous decrease as the populations approached the limits of responses (Supplementary Figure 6). Island PS demonstrated lesser rates of inbreeding compared to island GS and WGS. RM design showed the least rates of inbreeding among the four MDs for BI, RB and FC migration policies (Supplementary Figure 7 and 8). CR design followed a pattern similar to HN or GM depending on the SM. FC migration policy demonstrated lesser rates of inbreeding compared to BI and RB policies. BI policy demonstrated the largest rates of inbreeding. GM design demonstrated rates of inbreeding that were intermediate between RM and HN/CR designs (Supplementary Figure 8). Rs_Var for island selection with FC migration policies were larger than that observed with NI populations, demonstrating larger efficiency of converting loss of genetic variance into gain. However, with FC policy, all MDs showed a similar pattern (Supplementary Figure 9). Whereas Rs_Var for island selection with BI and RB policies were comparable to that of global selection with PS and GS, except for GM design, which showed larger Rs_Var after 10-20 cycles of selection (Supplementary Figure 10).

#### 4.4.1 Diversity within and among islands

The average within island genotypic variance decreased towards zero through forty cycles of selection, whereas global and inter-island genotypic variance increased before becoming limited. The rates of decrease in average within island genotypic variance was influenced by SM, MD and MP. Both GS and WGS demonstrated similar patterns of lost genotypic variance within islands and rates of loss with both were faster than PS (Figure 5). The HN mating design demonstrated the fastest loss of within island genotypic variance followed by RM, CR and GM designs. The FC migration policy provided the slowest loss of within island genotypic variance followed by RB and BI migration policies (Figure 5). Notice, however, an oscillatory pattern in which within island genotypic variance increased with every migration event and decreased from selection in cycles where there were no migrants. For both the within island genotypic variance and the expected heterozygosity the magnitudes of oscillations dampened towards zero after 20-30 cycles of selection except for the GM mating designs coupled with BI and GM migration policies (designated IM-GM-BI and IM-GM-RB respectively in Figure 5). The amplitude of increased genetic variance due to migration was greater for RB and BI migration policies with large spikes after 25-30 cycles of selection, while the amplitudes were smaller with the FC migration policies (Figure 5).

The largest values for inter-island genotypic variance were obtained with the RM mating design and BI and RB migration policies followed by CR and HN designs with BI and RB migration policies (Figure 7). Whereas, the FC migration policies demonstrated the smallest increases in inter-island genotypic variance through 40 cycles of selection (Figure 7). Recall that the FC migration policies provide the greatest migration rates among islands. Global genotypic variance in FI populations increased due to increase in inter-island genotypic variance. The BI migration policies demonstrated the largest global genetic variance for RM, HN and CR mating designs followed by the RB migration policies. Genomic mating under BI and RB migration policies provided intermediate rates of increasing global genotypic variance while the FC migration policy showed the least increase in global genotypic variance when coupled with the HN, CR, RM and GM mating designs (Figure 7).

**Figure 7.**
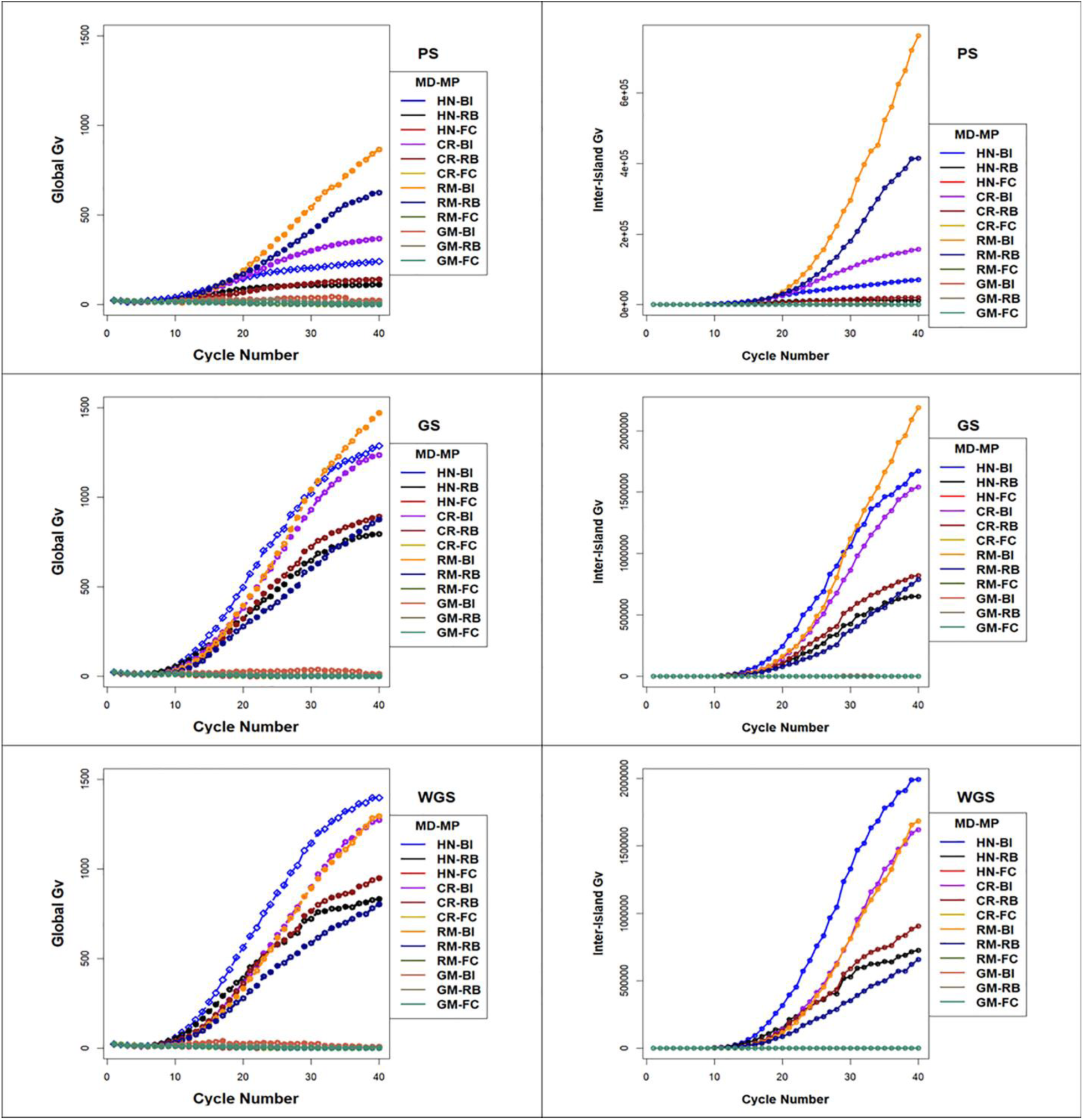
Global and Inter-Island Genotypic Variance in Island Selection i) **G**lobal genotypic variance (Left Panel), and ii) Inter-island genetic variance (Right Panel) for PS (top), GS (middle) and WGS (bottom) for the four mating designs including HN, CR, RM and GM methods and three migration policies including BI, RB and FC for 400 simulated QTL and 0.7 H. Genotypic variance is standardized to the average genotypic variance in founder populations in cycle ‘0’. GP models are updated every cycle in GS and WGS using training sets with data from all prior cycles of selection. PS- Phenotypic Selection, GS- Genomic Selection, WGS -Weighted Genomic Selection. Mating Design: HN (Hub Network), CR (Chain rule), and RM (Random Mating), GM (Genomic Mating) method. Migration policies in the island models consisted of bidirectional exchange of two immigrants and emigrants every other cycle of selection. Migration policies include BI- “Best Island”, RB- “Random Best”, and FC- “Fully Connected”

Within the classes of migration policies, the migration frequency had significant influence on rates and limits of responses across most combinations of selection methods, mating designs and migration policies, while numbers of migrants significantly affected responses in only for a few combinations of factors. Both rates and limits of response decreased with fewer migrants for the HN mating design. For the RM design, exchange of migrants among FI’s once in every three cycles provide the greatest genotypic values at the limits compared to responses with more frequent exchange. Migration size and migration direction had no significant effect on limits to selection responses (data available on request).

### 4.5 Tradeoffs between short-term and long-term gains from recurrent selection

There were 12 combinations of selection methods and mating designs applied to NI populations and 48 combinations of selection methods, mating designs and migration policies applied to FI populations. From among the 60 methods, genomic selection using a ridge regression model followed by a hub model mating design in NI populations and weighted genomic selection followed by crosses using the chain rule in NI populations (respectively designated NI-GS-HN and NI-WGS-CR in Table 2 and Figure 4) demonstrated the greatest responses in the first 20 and last 20 cycles respectively. However, if the objective for genetic improvement is to maximize gain in the first 5, 10, 30 or 40 cycles, other combinations of the factors are needed to achieve the objective. If the breeding objective is to maximize rates of genetic improvement in five to ten cycles of recurrent selection then there are two best options: 1. Genomic selection using RRBLUP values followed by a hub model mating design in FI populations with fully connected migration policies, or 2. Genomic selection using RRBLUP values followed by a genomic mating design in FI populations with fully connected migration policies (respectively designated as IM-GS-HN-FC and IM-GS-GM-FC in Table 2). If the objectives are to maximize both short-term and long-term gains then the best solution was obtained by selecting with RRBLUP values followed by a genomic mating in FI populations and applying a fully connected migration policy (designated IM-GS-GM-FC in Table 2). Among the combinations applied on NI populations, weighted genomic selection followed by the CR mating design or RM resulted in largest long-term gains, while selection using RRBLUP values followed by a HN mating design provided the greatest short-term gains. Indeed, both GS and WGS demonstrated greater long-term responses than phenotypic selection in both NI and FI populations.

**Table 2.**
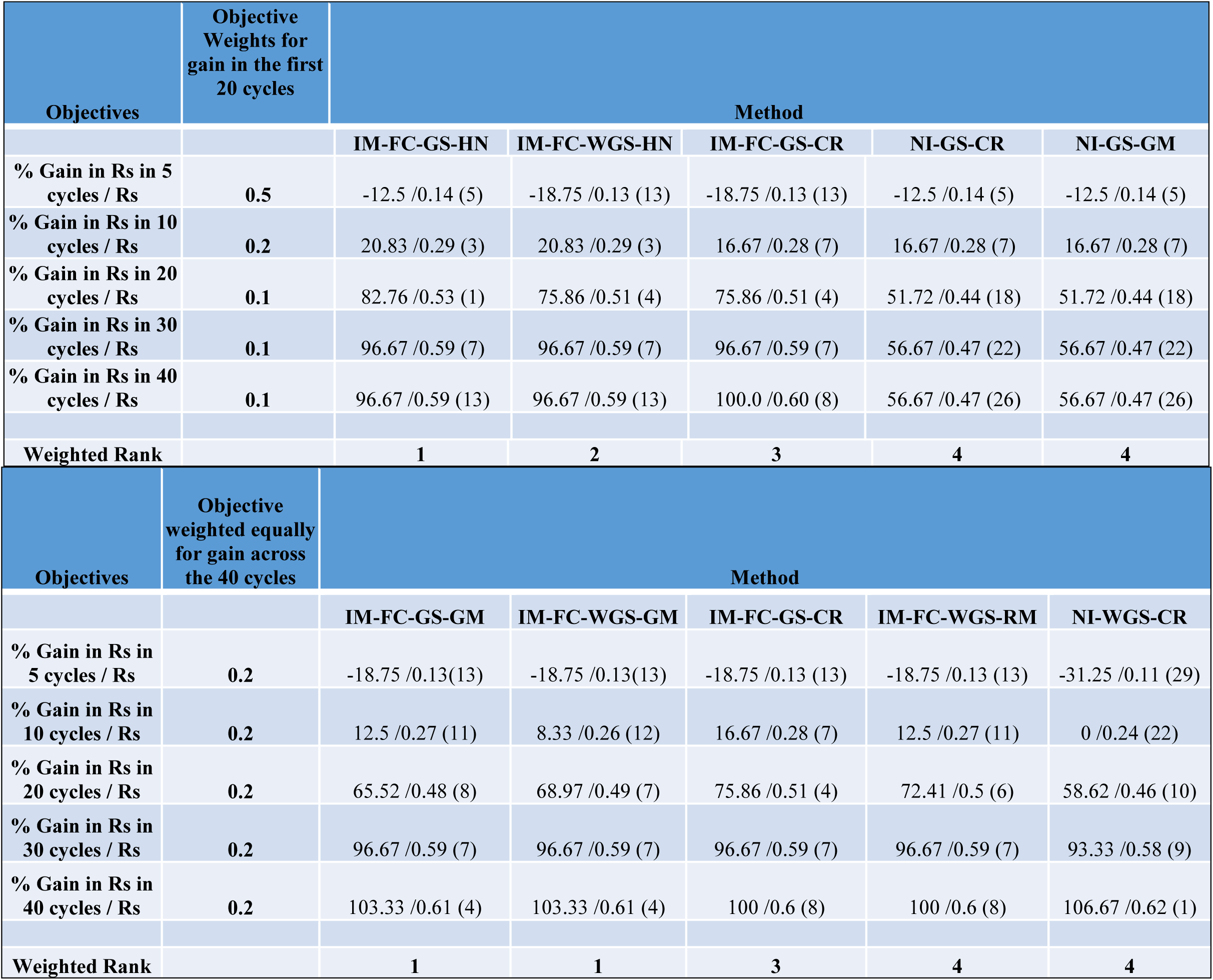
Trade-off Table for Strategies. Tradeoff table to support decision for selecting the best strategy to achieve possible objectives including maximum gain in 5, 10, 20, 30 and 40 cycles of recurrent selection. The methods are ranked for each of the objectives based on percent gain in genetic response relative to responses from PS on non-isolated families using a hub network mating design (NI-PS-HN). The relative gain relative to NI-PS-HN and the absolute genotypic response values for each of the methods are provided along with the ranking of the method for the specific objective within parenthesis. Three objective weights are provided to define the relative importance of the objectives: i) the weighted rank of methods are estimated with more emphasis on the first 20 cycles (top), ii) the weighted rank of methods are estimated with equal emphasis on the first and last 20 cycles (bottom). The best five methods among the 60 methods for each of the weighted objectives are presented. The simulations are provided for 400 simulated QTL responsible for 70% of phenotypic variability. Migration policies include “Discrete Selection”, “Best Island”, “Random Best”, and “Fully Connected”. Other migration factors correspond to migration frequency -2, migration direction -2 (bi-directional), and migration size - 2. Selection methods include PS-Phenotypic Selection, GS- Genomic Selection, and WGS-Weighted Genomic Selection. Mating Design includes HN (Hub Network), CR (Chain rule), RM- Random Mating, GM- Genomic Mating method.

## 5. Discussion

### 5.1 Significance

The challenge of finding optimal trade-offs among competing genetic improvement objectives has usually been approached by combining selection and crossing in a single step without consideration of population structure (Akdemir & Sanchez 2016; Beukelaer et al. 2017; Akdemir et al. 2019; Allier et al. 2019 a, b, 2020; Ramasubramanian and Beavis 2020). Akdemir & Sanchez (2016) combined selection and mating in their GM method. Beukelaer et al (2017) used weighted selection indices to maximize gain while retaining a threshold level of diversity. Among the three diversity measures they tested, indices that incorporate diversity measures to minimize loss of rare favorable alleles and minimize heterozygosity resulted in responses that were greater than WGS with truncation selection. Including diversity measures in a set offered advantage over truncation selection, as selected mate pairs retained rare favorable alleles better than WGS coupled with random mating. Allier et al (2019 a, b) included the impact of within-family selection to maximize genetic gain while minimizing loss of genetic variance, but they did not consider migration among families. And (Ramasubramanian and Beavis 2020) investigated GS methods for genetic improvement of soybean, but only considered the HN mating design applied to populations without regard to family affiliation. Herein we approached the challenge by disentangling breeding decisions into four distinct groups: 1) organization of the breeding population, 2) selection methods, 3) mating designs and 4) migration policies. Each of these were divided into possible alternatives within each group and treated as independent factors in orthogonal treatment combinations.

As with our previous investigation we found that the fastest rates of genetic improvement resulted when GS followed by the HN mating design is applied among all lines regardless of their family affiliation. When combined, these three decisions have reinforcing effects on responses to selection. At the other extreme, when WGS is applied to populations organized as family islands followed by either CR or RM the tendency of all three to retain genetic diversity reinforce each other resulting in the largest genotypic values, but is not achieved until the later cycles of selection. Because the slopes of the curves resulting from WGS and PS at 40 cycles are still positive, it is possible that both selection methods could continue to produce greater genetic potential with more cycles of selection. In previous comparative studies, WGS produced long-term responses that are similar to methods such as Optimal Contribution Selection (OCS) and Expected Maximum – Haploid Value (EM-HPV) (Daetwyler et al. 2015; Muller et al. 2018). Herein when we applied WGS to lines regardless of family affiliation and followed selection by identifying optimal mate pairs using GM the genotypic values at the limits to response were greater than the genotypic values obtained with PS or GS followed by GM. This combination also retained the largest values for heterozygosity and favorable alleles across more cycles. However, the differences between responses to GS and WGS followed by GM were not significant when applied to lines organized into family islands.

Between the extreme response curves it was also possible to find many response curves with intermediate trade-offs between the objectives. For example applying WGS to lines that were not organized into islands followed by HN provided greater response rates than other combinations of factors involving WGS. Selection among lines organized into FIs resulted in responses that were larger or comparable to responses from NI populations for only a limited number of combinations of mating design (GM) and migration policies (RB and FC). This may be due to the small numbers of related lines on each island (20X smaller than the NI population). With such a small number selection can deplete all the genetic variance within the first 10 -15 cycles as demonstrated in discrete selection. When there is no migration, which is the major source of new genetic variability, the populations realized only 10-15 % of maximum potential in the founder populations even while optimizing for sustainable gain using the GM method. A relaxed selection intensity, where the top 20% of the lines in each island are selected can sustain responses for longer cycles as demonstrated in non-isolated and island selection with migration (Supplementary Figure 12).

As expected, even with small numbers of lines per island migration had a positive impact on the outcomes. It is known that intermediate levels of migration rate result in optimal tradeoffs between gain and diversity (Skolicki 2007 a, b: Obolski et al. 2017). However, the range of intermediate parameter values depend on the specific context and population genetic parameters. In our study, responses in FIs were larger than selection responses in NI only when migration events happened every cycle or once in two cycles. When migration event happened only once in 3 cycles of selection, the rates of responses in the early cycles were very low resulting in much lower genotypic values as responses to selection approached limits. Migration size and direction didn’t have any significant impact on response within the small range of parameter values we tested for migration size and direction.

Also, we retained the best line within island during migration events and replaced the second best line in the ranked list of selected lines with the immigrant for the BI and RB policies. Whereas, for the FC policy, lines ranked from 2-6 are replaced. This replacement policy allows crossing between lines that are best within islands and immigrant lines from islands with high genotypic value resulting in high rates of response within islands. However, other policies that replace lines with low genotypic value with high genotypic values from immigrant islands will reduce genetic diversity within islands and result in different outcomes compared to the policy we’ve implemented.

None-the-less, we found a very good tradeoff among the competing objectives. If a GS was applied to lines on FC islands and the selected lines were mated according to the pareto-optimal crosses identified using GM, then the combination preserved genetic variance for long-term gain with little penalty relative to the realized rates of improvement in early cycles by GS and the HN mating design. In summary, motivated by Akdemir and Sanchez (2016) and Yabe et al (2016), we demonstrate that it is possible to design breeding strategies to produce desired outcomes between the extremes of maximizing the rate of genetic improvement and maximizing the genetic potential of the population.

### 5.2 Interpretations

The results can be interpreted from other perspectives to provide alternative insights. First genetic improvement can be viewed as single or multiple connected search strategies in genotypic space (Podlich and Cooper 1999; Cooper et al. 2002; Cooper et al. 2014). The single search strategy, a.k.a. global, corresponds to selection of lines in NI populations. The multiple connected search strategy, a.k.a. local, corresponds to selection of lines in multiple domains with infrequent exchange of lines. In addition to the perspective of global or local search strategies, selection can be viewed as cooperation vs. competition and exploitation vs. exploration. Thus, by tuning parameters that control relative levels of cooperation or exploration in global or local search strategies, it is possible to adjust the adaptive landscape. Genotypes co-operate when they contribute to other genotypes’ fitness values, whereas they compete when they reduce the fitness values of other genotypes thereby reducing their contribution to future generations. Intermediate levels of cooperation often accelerated shifts in adaptive peaks for bi-locus genetic models (Whitley 1999; Skolicki 2007; Obolski et al. 2017). For global selection, selection methods and mating designs control contributions of genotypes within populations thereby controlling the level of cooperation. From this perspective, GS promotes competition, while PS and WGS emphasize cooperation on the adaptive landscape. By weighting rare favorable alleles, WGS promotes cooperation and effectively retains more of the genotypic potential of the founder’s fitness landscape. Among mating designs, the HN used by most plant breeders, promotes competition over cooperation, whereas the CR and RM designs promote co-operation. The GM design provides a balance between cooperation and competition. For island selection methods, a FC migration policy provides the best balance between cooperation and competition. Further work is needed to identify optimal migration rules.

The concepts of exploitation and exploration are commonly used in EAs. In general, exploitation refers to processes such as selection that result in beneficial solutions, whereas exploration allows searches for solutions in new domains. In the breeding context, exploration maintains diversity (Goldberg 1989; Goldberg and Deb 1992; Whitley 1999; Skolicki 2007 a, b; Crepinsek et al. 2013). Because GS provides faster rates of genetic improvement than PS and WGS it is reasonable to interpret GS results as rapid exploitation, whereas PS and WGS allow exploration of new solutions, primarily through additional recombination opportunities. The HN mating design drives the populations to exploit resources for immediate gains, whereas CR and RM mating designs provide opportunities for the population to explore more of the fitness landscape. The GM mating design provides an opportunity to choose relative importance of exploitation and exploration. By treating population organization, selection methods and mating designs as orthogonal factors we were able to blur the boundary between exploitation and exploration with combinations of factors that mixed exploitation and exploration activities in distinct phases and simultaneously.

From the perspective of tradeoffs between exploration and exploitation, selection among and within islands enables exploration of diverse domains resulting in greater probabilities of finding solutions with greater genotypic values (higher peaks or limits at convergence), whereas global selection across a NI population of lines tends to get trapped in sub-optimal peaks or local optima (Cantu-Paz 2000; Skolicki 2007 a, b; Luque 2011; Crepinsek et al. 2013). In our study, this occurred with the GM design applied to the NI populations, where the crossing process within islands are optimized at a local level. In some implementations of island model selection, both the global and local states of islands are assessed every cycle of selection and a centralized global agent makes decisions to reach optimization objectives. We do not know whether such a bi-level optimization method will result in greater genotypic values before approaching limits to responses from selection.

### 5.3 Future research

By framing breeding strategies as combinations of population structure, selection methods, mating designs and migration policies we illustrated the potential of the approach for a small arbitrary soybean genetic improvement project. We did not consider the relative emphasis of objectives and constraints for any specific genetic improvement project. Consider first the structure of breeding populations. We compared a NI structure of lines with FI’s created by individual crosses among the founders and then we selected within and among islands according to the same criteria. This might make sense within a single soybean genetic improvement project for lines adapted to MZ’s II and III.

There are six public soybean genetic improvement projects adapted MZ’s II and III. There are likewise about the same number of commercial soybean genetic improvement projects in the same MZs. All of these projects began at different times and were initiated with unique, albeit overlapping, germplasm resources (Mikel et al. 2010). While all of the projects select lines with greater genotypic values for yield, the yield values are obtained from different, albeit overlapping environments.

From the perspective of soybean genetic improvement across regions within MZ’s II and III, each genetic improvement project can be represented as an island where genotypes are exchanged among project islands based on annual evaluations in uniform regional trials and according to legal licensing rules. In practice project islands exchange only the best performing lines adapted to similar environmental conditions. None-the-less, soybean breeders will maintain useful genetic variability by exchanging lines among island projects. An advantage of island selection is that diversity among islands increases with selection, even when within island diversity decreases. Eventually, beyond 40 cycles of recurrent selection, genetic variability among islands will decrease as genetic variability among islands is lost to selection.

Future investigations of breeding strategies to optimize tradeoffs between rates of genetic gains and retention of useful genetic variance in soybean adapted to MZ’s II and III should consider population structures within island projects that more accurately reflect those that currently exist. Also, future investigations should simulate genetic architectures with genotype x environment effects. It is well known that a line adapted to one environment may not perform well in other environments, and it is possible to define fitness values so that they include environmental effects. Third, future investigations should consider a broader set of migration rules and policies. The FC migration policy is considered the upper bound of island models as all islands are connected to every other island with maximum migration rates among islands. While our results indicate that this policy provided the results we don’t think it will provide the best results for genetic architectures with genotype by environment interaction effects.

Fourth, we need to recognize islands in time because every cycle of selection discards useful genetic variability. A soybean germplasm resource project was set up (Mikel et al. 2010) to recover useful genetic variability lost during domestication of soybean (Nelson 2011). Rather than trying to build long bridges to islands located in the distant past, our results suggest that there should be a large amount of useful genetic variability that was discarded in the first few cycles of modern soybean breeding. For that matter, until response to selection reaches the half-life for the population, large amounts of useful genetic variability can probably be recovered from islands represented by recent cycles of discarded lines. These conjectures should be preceded by simulations to determine the potential benefit and costs associated with sampling lines in recently discarded islands.

Fifth, it should be clear that a predefined mating design does not take advantage of opportunities created by each cycle of progeny to optimize outcomes according to most project objectives. Thus, there continues to be a need for algorithms that efficiently and effectively identify crosses from among genotypes produced by each cycle of selection. It is tempting to adopt and investigate all EA strategies. However, only a subset are relevant to the practice of plant breeding (Hagan et al. 2012). For example, mutation and recombination rates can be controlled in a computational EA, whereas plant breeders cannot regulate these with current practices. None-the-less there are many opportunities for cross-disciplinary research between EAs and plant breeding. There is large body of literature concerning the properties of EAs and factors and strategies that affect convergence rates and quality of solutions (Goldberg 1989; Goldberg and Deb 1992; Whitley 1999; Skolicki 2007 a, b; Crepinsek et al. 2013; Obolski et al. 2017) and working with computational scientists should reveal novel methods to maximize the genetic potential of a breeding population in a minimum number of cycles.

Akdemir and Sanchez (2016) proposed only one of many possible GA’s to identify pareto-optimal solution pairs. An approach introduced by Gaur and Deb (2016) and Mittal et al. (2020) would provide pareto-optimal solutions using statistical methods such as clustering and machine learning. The statistical knowledge can be used to improve the search for optimal solutions and establish several cycles of optimization. Conceptually, unveiling any relationship among pareto-optimal pairs in a genotypic space is likely to provide new knowledge regarding the characteristics of such complementary pairs. Also, by modeling responses with a first order recurrence equation or a non-linear mixed effects model to predict the half-life and asymptotic limits of selection have potential to improve the efficiency of genetic algorithms by providing repair operators to alter the trajectory of population evolution towards the desired optimal trade-offs.

Last, consider the challenge of stating explicit relative emphasis on objectives and definition of constraints for any specific genetic improvement project. As noted previously, this challenge exists because it requires assigning economic and agronomic value of short term genetic gains vs. the forecasted value of useful genetic variants that may be discarded each cycle of selection. As a thought experiment note that the trade-off objectives can be reduced to a single ‘grand’ objective of creating a genotype (line) with the genotypic value equal to the full genetic potential of the founders in a single cycle. For a genetic architecture consisting of two alleles at a single locus, achieving the single grand objective is trivial. Also it is possible to imagine that the grand objective can be achieved for a complex genetic architecture with infinite resources. Clearly, given genetic architectures of complex traits and resource constraints there are no feasible solutions to the grand objective, but it is a useful reference to serve as the goal.

## Acknowledgement

Funding for this research was provided by the Department of Agronomy, Iowa State University, the North Central Soybean Research Program and an NSF grant (1830478). Supplemental funding for large scale computing was enabled by the Extreme Science and Engineering Discovery Environment (XSEDE), which is supported by National Science Foundation. XSEDE resources consisted of research allocations (DMS180041, DMS190003, DMS190015 & DMS190018) on PSC-Bridges Large Memory nodes for the simulations involving island model and genomic mating simulations. We want to thank Dr. Deniz Akdemir for discussions on implementing ‘genomic mating’ and Dr. Lizhi Wang for efficient programs to simulate meiosis. We also want to thank Dr. Alencar Xavier for sharing an efficient expectation maximization method for fitting ridge regression GP models.

## Notes

http://gfspopgen.agron.iastate.edu/SoyNAMSelectionMethods_v2_2020.html

### Competing Interest Statement

The authors have declared no competing interest.

