## Supplemental Figures for "Strategies to assure optimal trade-offs among competing objectives for genetic improvement of soybean"

**Supplementary Figures**

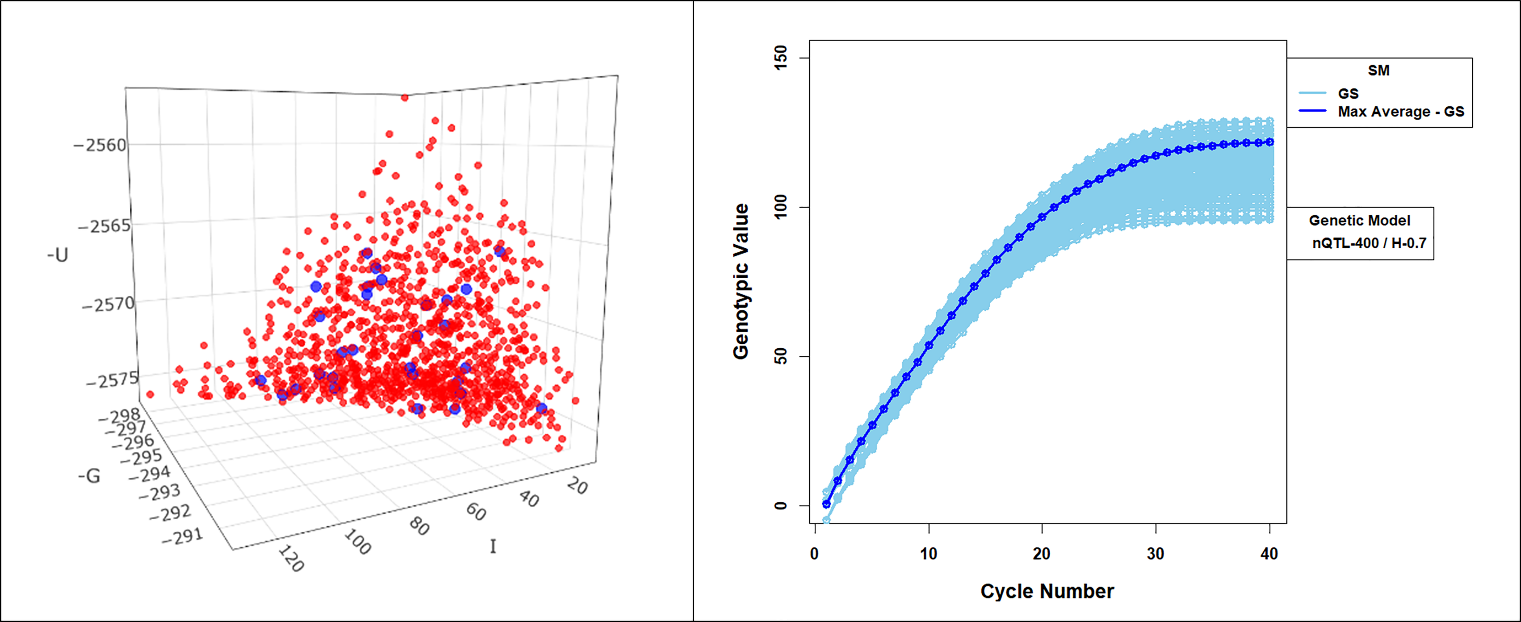

**Supplementary Figure1** The left panel is a plot of pareto-optimal solutions creating a frontier surface for values of genetic gain (-G), Inbreeding (I) and Usefulness (-U). For this particular plot, the GM method is applied to RILs selected in one of the family islands using genomic selection (GS). Each point is an optimal solution corresponding to a set of 10 pairs of RILs selected as parental lines within one island. A similar set of pareto-optimal solutions are selected for all 20 family islands. Out of all the pareto-optimal solutions (red circles) on the frontier surface, 30 (blue circles) consisting of 10 optimal solutions with high G, 10 optimal solutions with low G and 10 optimal solutions with median G are selected. The solution pairs in each of the 30 selected sets are crossed and the recurrent selection cycle is iterated for 40 cycles for each of the 30 sets.

The right panel is a plot of genotypic values, when the selected set of 30 solutions are crossed and the process of selection and crossing is iterated for 40 cycles of recurrent selection for all 20 family islands. The thick blue curves represent parameters with maximum limits of response for GS. The genotypic values represent responses in recurrent GS with 400 QTL responsible for 70% of phenotypic variability in the founding set of RILs and top 10% selected fraction with bi-directional migrations of two migrants every other cycle with FC (“Fully Connected”) migration policy.

**`**

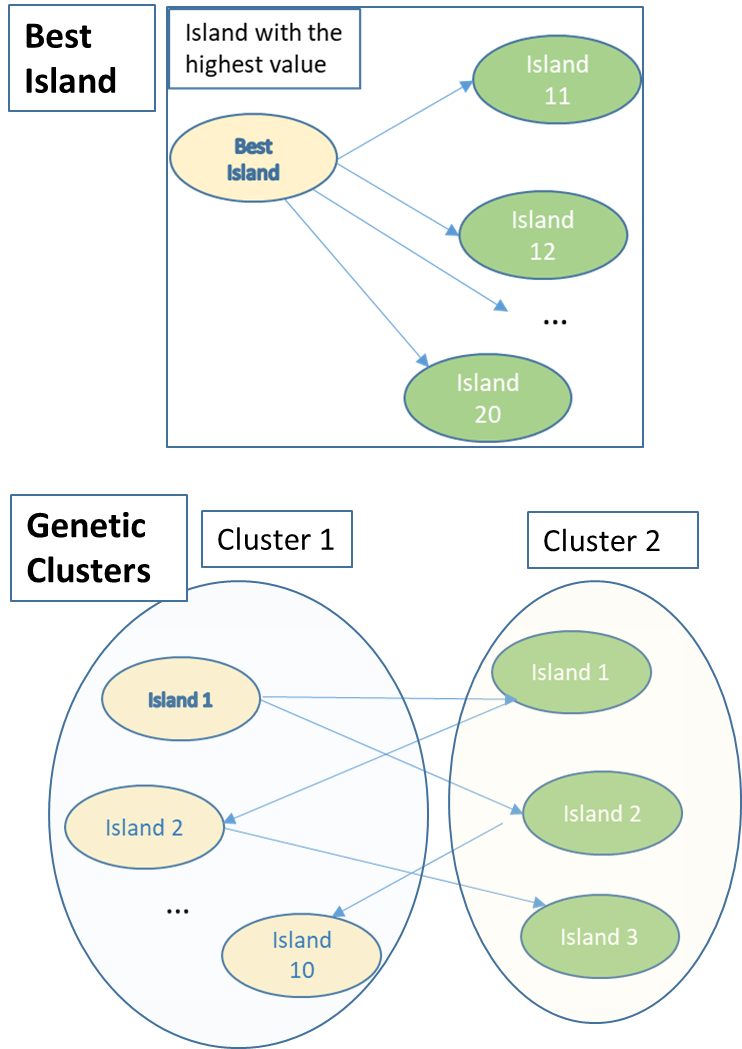

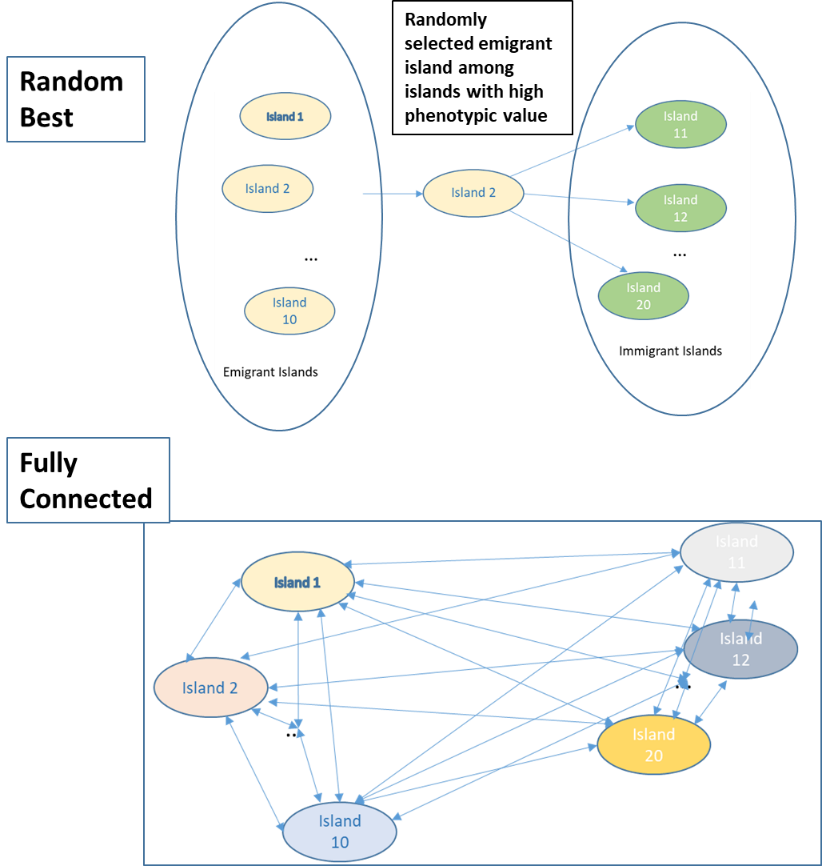

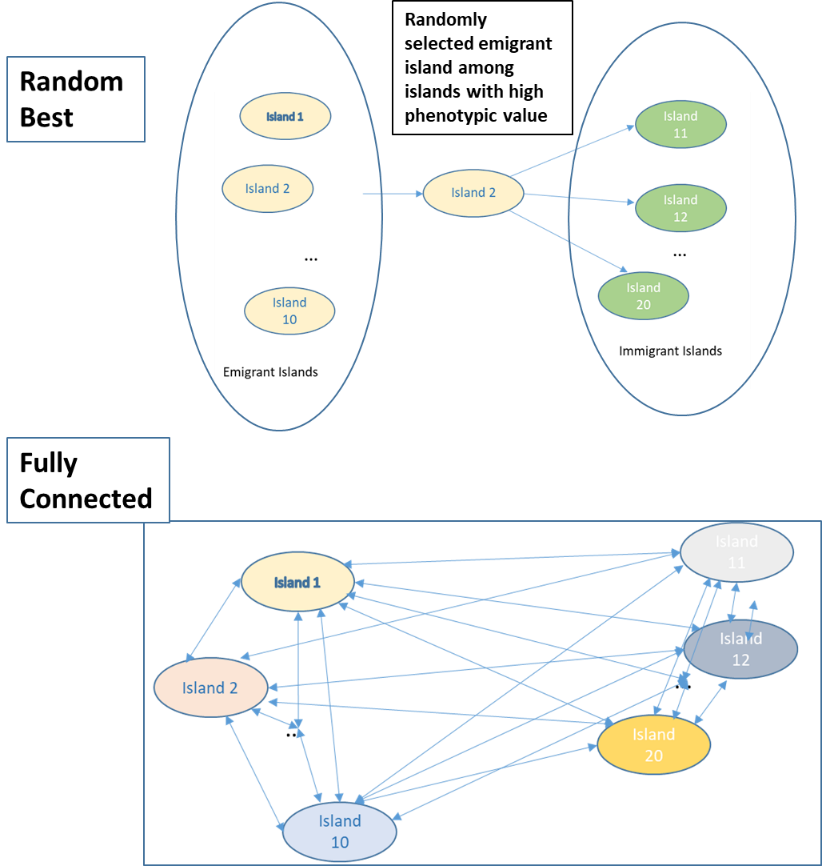

**Supplementary Figure2.** Representations of Island Topology and Migration Policies**: “**Best Island” (top-left) designates that emigrant lines are selected from the island with the largest genotypic value. Emigrants migrate to no more than 10 islands. If migration direction is bi-directional, the emigrant island also receives immigrants; ii) “Random Best” (top-right), denotes that an emigrant island is selected randomly from a set of 10 islands with large genotypic values. The emigration pattern is similar to “Best Island” policy iii) A “Fully connected” island topology is represented in the bottom panel and is defined as every island is connected to every other island. In the fully connected topology lines migrate from emigrant islands with high values to randomly selected immigrant islands.

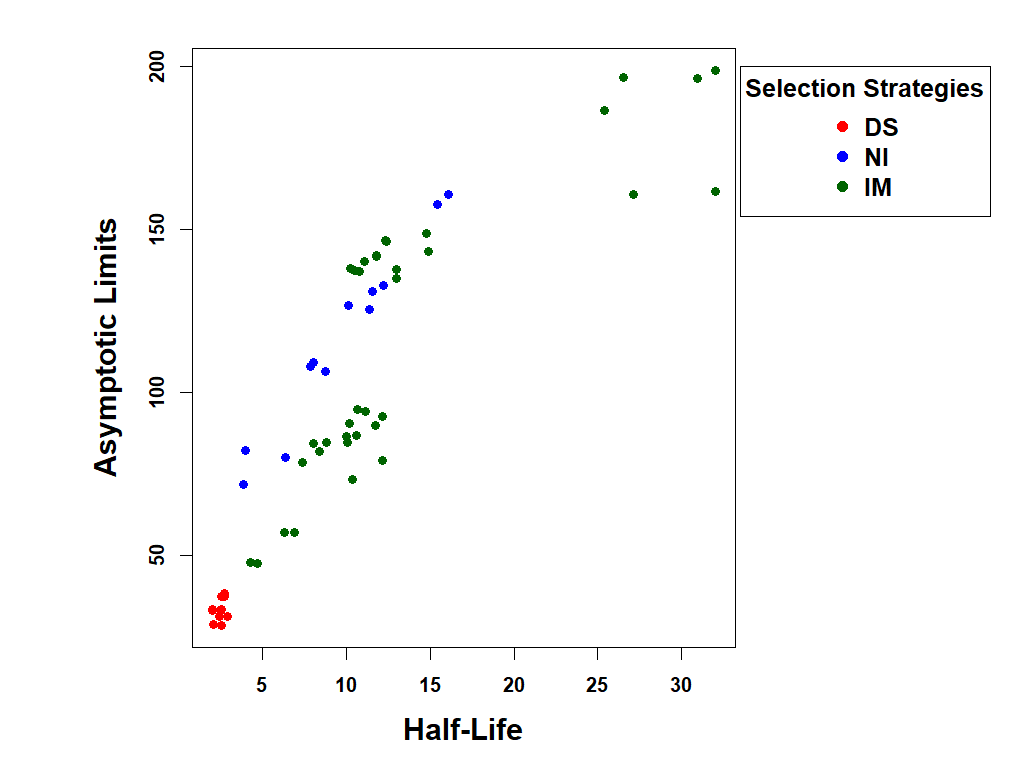

**Supplementary Figure3. Asymptotic limits and Half-life of Recurrent Selection for the 60 methods**. Half-life is plotted on the x-axis and asymptotic limits on the y-axis. The cluster of red points correspond to discrete selection (DS) methods. The blue and green points correspond to non-isolated (NI) and island selection (IM) methods. The cluster of green points in the top-right corner with half-life above 25 and asymptotic limits above 160 correspond to island selection with GM method combined with Best Island and Random Best migration policies (IM-GM-BI & IM-GM-RB with PS, GS and WGS). These points are likely to be over-estimates, as the genotypic values for 40 cycles of selection fall on the linear region of the response curve. Simulations of additional cycles of selection are likely to provide more accurate estimates for these methods with larger half-life and asymptotic limits compared to all other methods, but lesser than the current estimates. All simulations use 400 simulated QTL responsible for 70% of phenotypic variability. Top 10% of the lines are selected from non-isolated and island populations as parental lines to be crossed using HN, CR, RM and GM mating designs. Selection Methods: PS-Phenotypic Selection, GS- Genomic Selection, WGS -Weighted Genomic Selection with Jannink weighting function. Mating Design: HN (Hub Network), CR (Chain rule), RM- Random Mating, GM- Genomic Mating method. Migration policies: “Discrete Selection” (DS), “Best Island” (BI), “Random Best” (RB), and “Fully Connected” (FC).

| **Mating Design** | | **Selection Policy** | **Selection Method** | **5** | **10** | | **20** | | **30** | | **40** | |
| --- | --- | --- | --- | --- | --- | --- | --- | --- | --- | --- | --- | --- |
| **Hub Network** | **Non-Isolated** | | **PS** | **0 (0.16)** | | **0 (0.24)** | | **0 (0.29)** | | **0 (0.3)** | | **0 (0.3)** |
|  |  |  | **GS** | **12.5 (0.18)** | | **20.83 (0.29)** | | **20.69 (0.35)** | | **13.33 (0.34)** | | **16.67 (0.35)** |
|  |  |  | **WGS** | **-25 (0.12)** | | **-8.33 (0.22)** | | **10.34 (0.32)** | | **13.33 (0.34)** | | **16.67 (0.35)** |
|  | **Discrete Selection** | | **PS** | **-50 (0.08)** | | **-58.33 (0.1)** | | **-65.52 (0.1)** | | **-66.67 (0.1)** | | **-66.67 (0.1)** |
|  |  |  | **GS** | **-43.75 (0.09)** | | **-54.17 (0.11)** | | **-62.07 (0.11)** | | **-63.33 (0.11)** | | **-63.33 (0.11)** |
|  |  |  | **WGS** | **-43.75 (0.09)** | | **-50 (0.12)** | | **-58.62 (0.12)** | | **-60 (0.12)** | | **-60 (0.12)** |
|  | **Best Island** | | **PS** | **-43.75 (0.09)** | | **-37.5 (0.15)** | | **-34.48 (0.19)** | | **-33.33 (0.2)** | | **-30 (0.21)** |
|  |  |  | **GS** | **-31.25 (0.11)** | | **-20.83 (0.19)** | | **0 (0.29)** | | **13.33 (0.34)** | | **20 (0.36)** |
|  |  |  | **WGS** | **-31.25 (0.11)** | | **-20.83 (0.19)** | | **3.45 (0.3)** | | **16.67 (0.35)** | | **23.33 (0.37)** |
|  | **Random Best** | | **PS** | **-37.5 (0.1)** | | **-33.33 (0.16)** | | **-34.48 (0.19)** | | **-33.33 (0.2)** | | **-33.33 (0.2)** |
|  |  |  | **GS** | **-25 (0.12)** | | **-16.67 (0.2)** | | **-3.45 (0.28)** | | **10 (0.33)** | | **16.67 (0.35)** |
|  |  |  | **WGS** | **-25 (0.12)** | | **-16.67 (0.2)** | | **0 (0.29)** | | **16.67 (0.35)** | | **20 (0.36)** |
|  | **Fully Connected** | | **PS** | **-31.25 (0.11)** | | **0 (0.24)** | | **48.28 (0.43)** | | **76.67 (0.53)** | | **86.67 (0.56)** |
|  |  |  | **GS** | **-12.5 (0.14)** | | **20.83 (0.29)** | | **82.76 (0.53)** | | **96.67 (0.59)** | | **96.67 (0.59)** |
|  |  |  | **WGS** | **-18.75 (0.13)** | | **20.83 (0.29)** | | **75.86 (0.51)** | | **96.67 (0.59)** | | **96.67 (0.59)** |
| **Chain Rule** | **Non-Isolated** | | **PS** | **-25 (0.12)** | | **4.17 (0.25)** | | **55.17 (0.45)** | | **76.67 (0.53)** | | **86.67 (0.56)** |
|  |  |  | **GS** | **-12.5 (0.14)** | | **16.67 (0.28)** | | **51.72 (0.44)** | | **56.67 (0.47)** | | **56.67 (0.47)** |
|  |  |  | **WGS** | **-31.25 (0.11)** | | **0 (0.24)** | | **58.62 (0.46)** | | **93.33 (0.58)** | | **106.67 (0.62)** |
|  | **Discrete Selection** | | **PS** | **-56.25 (0.07)** | | **-54.17 (0.11)** | | **-58.62 (0.12)** | | **-60 (0.12)** | | **-60 (0.12)** |
|  |  |  | **GS** | **-43.75 (0.09)** | | **-50 (0.12)** | | **-58.62 (0.12)** | | **-60 (0.12)** | | **-60 (0.12)** |
|  |  |  | **WGS** | **-43.75 (0.09)** | | **-50 (0.12)** | | **-58.62 (0.12)** | | **-60 (0.12)** | | **-60 (0.12)** |
|  | **Best Island** | | **PS** | **-43.75 (0.09)** | | **-37.5 (0.15)** | | **-24.14 (0.22)** | | **-20 (0.24)** | | **-16.67 (0.25)** |
|  |  |  | **GS** | **-37.5 (0.1)** | | **-25 (0.18)** | | **-3.45 (0.28)** | | **16.67 (0.35)** | | **26.67 (0.38)** |
|  |  |  | **WGS** | **-37.5 (0.1)** | | **-25 (0.18)** | | **-3.45 (0.28)** | | **13.33 (0.34)** | | **23.33 (0.37)** |
|  | **Random Best** | | **PS** | **-43.75 (0.09)** | | **-33.33 (0.16)** | | **-20.69 (0.23)** | | **-20 (0.24)** | | **-16.67 (0.25)** |
|  |  |  | **GS** | **-31.25 (0.11)** | | **-20.83 (0.19)** | | **6.9 (0.31)** | | **26.67 (0.38)** | | **33.33 (0.4)** |
|  |  |  | **WGS** | **-37.5 (0.1)** | | **-20.83 (0.19)** | | **3.45 (0.3)** | | **26.67 (0.38)** | | **33.33 (0.4)** |
|  | **Fully Connected** | | **PS** | **-31.25 (0.11)** | | **4.17 (0.25)** | | **55.17 (0.45)** | | **80 (0.54)** | | **90 (0.57)** |
|  |  |  | **GS** | **-18.75 (0.13)** | | **16.67 (0.28)** | | **75.86 (0.51)** | | **96.67 (0.59)** | | **100 (0.6)** |
|  |  |  | **WGS** | **-18.75 (0.13)** | | **16.67 (0.28)** | | **75.86 (0.51)** | | **93.33 (0.58)** | | **96.67 (0.59)** |
| **Random Mating** | **Non-Isolated** | | **PS** | **-25 (0.12)** | | **0 (0.24)** | | **51.72 (0.44)** | | **76.67 (0.53)** | | **83.33 (0.55)** |
|  |  |  | **GS** | **-18.75 (0.13)** | | **12.5 (0.27)** | | **62.07 (0.47)** | | **80 (0.54)** | | **83.33 (0.55)** |
|  |  |  | **WGS** | **-31.25 (0.11)** | | **0 (0.24)** | | **51.72 (0.44)** | | **86.67 (0.56)** | | **103.33 (0.61)** |
|  | **Discrete Selection** | | **PS** | **-56.25 (0.07)** | | **-58.33 (0.1)** | | **-62.07 (0.11)** | | **-63.33 (0.11)** | | **-63.33 (0.11)** |
|  |  |  | **GS** | **-50 (0.08)** | | **-54.17 (0.11)** | | **-62.07 (0.11)** | | **-63.33 (0.11)** | | **-63.33 (0.11)** |
|  |  |  | **WGS** | **-50 (0.08)** | | **-54.17 (0.11)** | | **-62.07 (0.11)** | | **-63.33 (0.11)** | | **-63.33 (0.11)** |
|  | **Best Island** | | **PS** | **-50 (0.08)** | | **-41.67 (0.14)** | | **-20.69 (0.23)** | | **-3.33 (0.29)** | | **10 (0.33)** |
|  |  |  | **GS** | **-37.5 (0.1)** | | **-29.17 (0.17)** | | **-3.45 (0.28)** | | **16.67 (0.35)** | | **26.67 (0.38)** |
|  |  |  | **WGS** | **-43.75 (0.09)** | | **-29.17 (0.17)** | | **-6.9 (0.27)** | | **16.67 (0.35)** | | **26.67 (0.38)** |
|  | **Random Best** | | **PS** | **-43.75 (0.09)** | | **-37.5 (0.15)** | | **-20.69 (0.23)** | | **-3.33 (0.29)** | | **6.67 (0.32)** |
|  |  |  | **GS** | **-37.5 (0.1)** | | **-20.83 (0.19)** | | **3.45 (0.3)** | | **20 (0.36)** | | **33.33 (0.4)** |
|  |  |  | **WGS** | **-37.5 (0.1)** | | **-25 (0.18)** | | **0 (0.29)** | | **13.33 (0.34)** | | **23.33 (0.37)** |
|  | **Fully Connected** | | **PS** | **-31.25 (0.11)** | | **0 (0.24)** | | **51.72 (0.44)** | | **83.33 (0.55)** | | **96.67 (0.59)** |
|  |  |  | **GS** | **-25 (0.12)** | | **12.5 (0.27)** | | **72.41 (0.5)** | | **96.67 (0.59)** | | **100 (0.6)** |
|  |  |  | **WGS** | **-18.75 (0.13)** | | **12.5 (0.27)** | | **72.41 (0.5)** | | **96.67 (0.59)** | | **100 (0.6)** |
| **Genomic Mating** | **Non-Isolated** | | **PS** | **-18.75 (0.13)** | | **4.17 (0.25)** | | **41.38 (0.41)** | | **53.33 (0.46)** | | **56.67 (0.47)** |
|  |  |  | **GS** | **-12.5 (0.14)** | | **16.67 (0.28)** | | **51.72 (0.44)** | | **56.67 (0.47)** | | **56.67 (0.47)** |
|  |  |  | **WGS** | **-25 (0.12)** | | **0 (0.24)** | | **51.72 (0.44)** | | **73.33 (0.52)** | | **76.67 (0.53)** |
|  | **Discrete Selection** | | **PS** | **-37.5 (0.1)** | | **-45.83 (0.13)** | | **-51.72 (0.14)** | | **-53.33 (0.14)** | | **-53.33 (0.14)** |
|  |  |  | **GS** | **-37.5 (0.1)** | | **-45.83 (0.13)** | | **-51.72 (0.14)** | | **-53.33 (0.14)** | | **-53.33 (0.14)** |
|  |  |  | **WGS** | **-37.5 (0.1)** | | **-41.67 (0.14)** | | **-51.72 (0.14)** | | **-53.33 (0.14)** | | **-53.33 (0.14)** |
|  | **Best Island** | | **PS** | **-50 (0.08)** | | **-41.67 (0.14)** | | **-13.79 (0.25)** | | **16.67 (0.35)** | | **46.67 (0.44)** |
|  |  |  | **GS** | **-37.5 (0.1)** | | **-29.17 (0.17)** | | **6.9 (0.31)** | | **46.67 (0.44)** | | **83.33 (0.55)** |
|  |  |  | **WGS** | **-37.5 (0.1)** | | **-25 (0.18)** | | **10.34 (0.32)** | | **50 (0.45)** | | **83.33 (0.55)** |
|  | **Random Best** | | **PS** | **-43.75 (0.09)** | | **-33.33 (0.16)** | | **0 (0.29)** | | **33.33 (0.4)** | | **60 (0.48)** |
|  |  |  | **GS** | **-37.5 (0.1)** | | **-20.83 (0.19)** | | **24.14 (0.36)** | | **66.67 (0.5)** | | **100 (0.6)** |
|  |  |  | **WGS** | **-37.5 (0.1)** | | **-20.83 (0.19)** | | **24.14 (0.36)** | | **66.67 (0.5)** | | **96.67 (0.59)** |
|  | **Fully Connected** | | **PS** | **-31.25 (0.11)** | | **-4.17 (0.23)** | | **41.38 (0.41)** | | **73.33 (0.52)** | | **90 (0.57)** |
|  |  |  | **GS** | **-18.75 (0.13)** | | **12.5 (0.27)** | | **65.52 (0.48)** | | **96.67 (0.59)** | | **103.33 (0.61)** |
|  |  |  | **WGS** | **-18.75 (0.13)** | | **8.33 (0.26)** | | **68.97 (0.49)** | | **96.67 (0.59)** | | **103.33 (0.61)** |

**Supplementary Figure4.** Heat map of relative genetic gain represented as a percentage of genetic response (Rs) relative to responses in NI-PS-HN in simulations for 400 simulated QTL responsible for 70% of phenotypic variability. Top 10% of the lines are selected from non-isolated and island populations as parental lines to be crossed using HN, CR, RM and GM mating designs. Selection Methods: PS-Phenotypic Selection, GS- Genomic Selection, WGS -Weighted Genomic Selection with Jannink weighting function. Mating Design: HN (Hub Network), CR (Chain rule), RM- Random Mating, GM- Genomic Mating method. Migration policies: “Discrete Selection” (DS), “Best Island” (BI), “Random Best” (RB), and “Fully Connected” (FC).

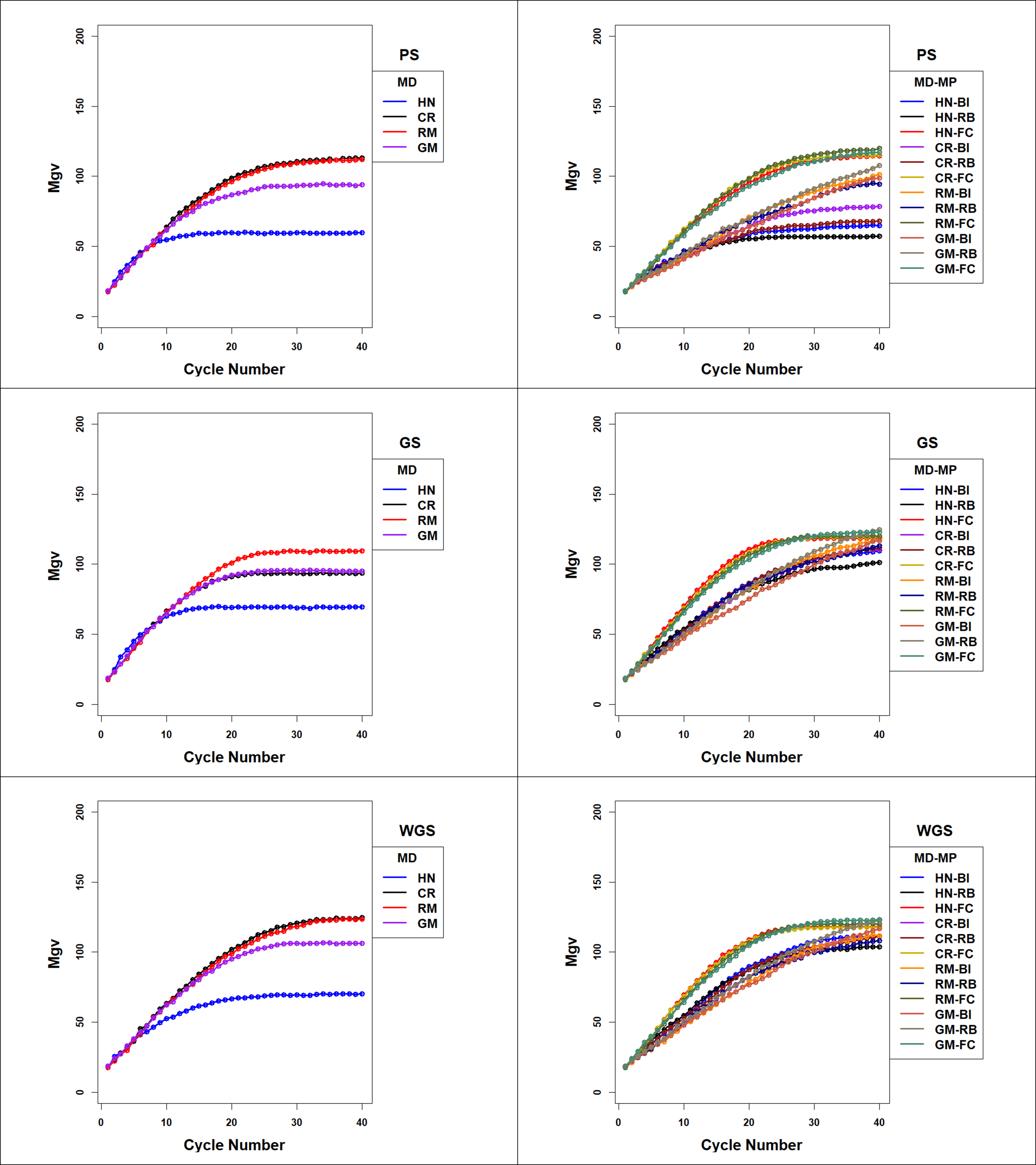

**Supplementary Figure 5.** Maximal Genotypic Values (Mgvs) across 40 cycles of recurrent selection on Non-isolated (left panels) and Island (right panels) populations, using Phenotypic Selection (PS-top panel), Genomic Selection (GS-middle panel) and Weighted Genomic Selection (WGS-bottom panel) and four mating designs: Hub-Network (HN), Chain Rule (CR), Random Mating (RM), and Genomic Mating (GM). Standardized genotypic responses are represented from a simulated genetic architecture consisting of 400 additive QTL uniformly distributed throughout the genome and responsible for 70% of phenotypic variability. Ten percent of lines are selected to be used in crosses in HN, CR, RM and GM designs. Migration policies include bi-directional migrations of two migrants every other cycle involving the Best Island (BI), Random Best (RB), and Fully Connected (FC) islands.

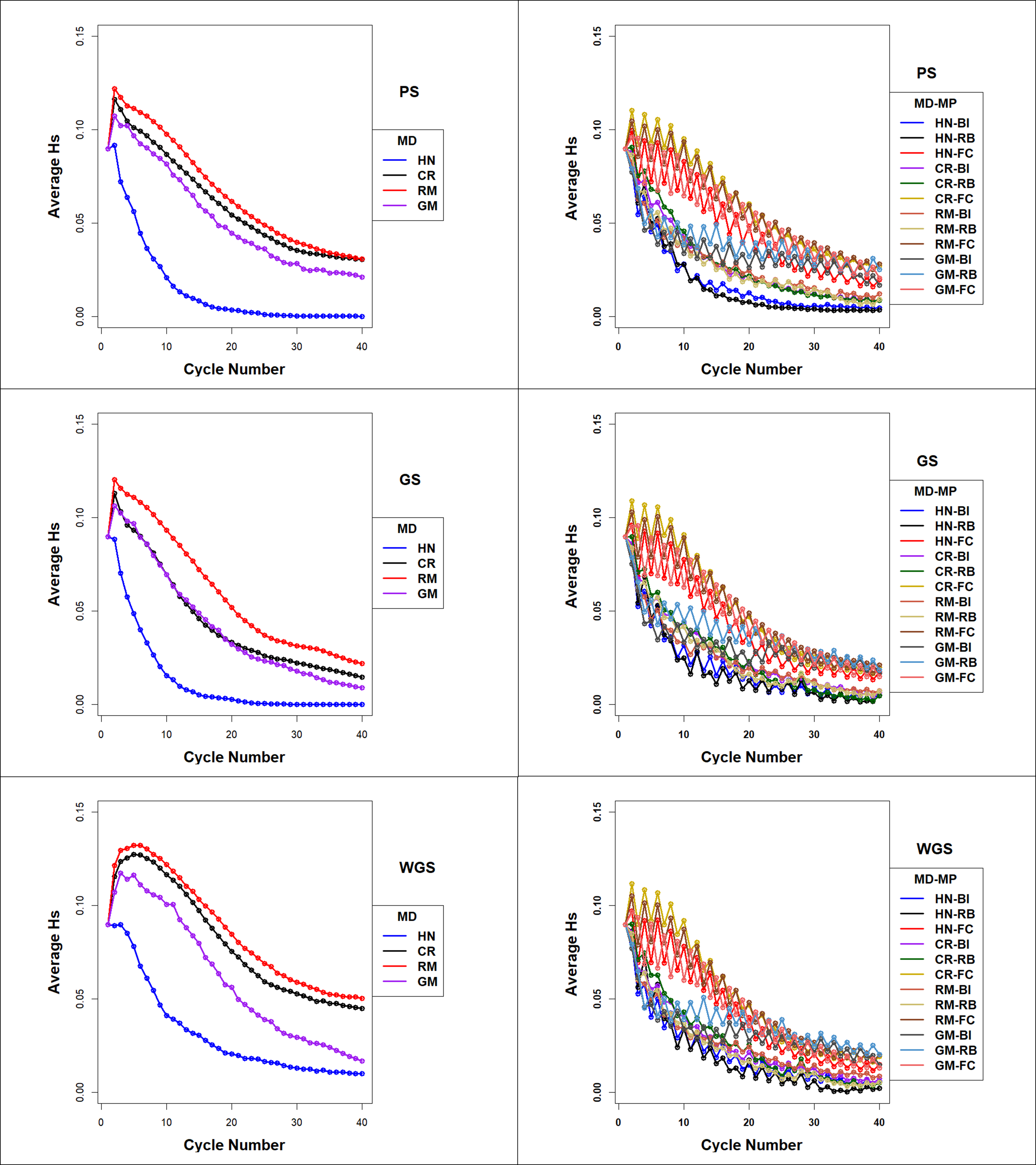

**Supplementary Figure 6** Average Expected Heterozygosity (Hs) in Non-isolated (NI) and Island populations organized as Island of families (FI). Hs in Non-isolated selection PS (top), GS (middle) and WGS (bottom) for the four mating designs including HN, CR, RM, and GM with a selection intensity of top 10% selected fraction. GP models are updated every cycle in GS and WGS, ii) Average Expected Heterozygosity PS (top), GS (middle) and WGS (bottom) with Jannink’s weighting function for the four mating designs including HN, CR, RM, and GM designs. GS models are updated every cycle. Selection Methods include PS-Phenotypic Selection, GS- Genomic Selection, and WGS -Weighted Genomic Selection with Jannink weighting function. Mating Designs include HN (Hub Network), CR (Chain rule), RM- Random Mating, GM- Genomic Mating method. Migration policies include bi-directional migrations of two migrants every other cycle involving the Best Island (BI), Random Best (RB), and Fully Connected (FC) islands.

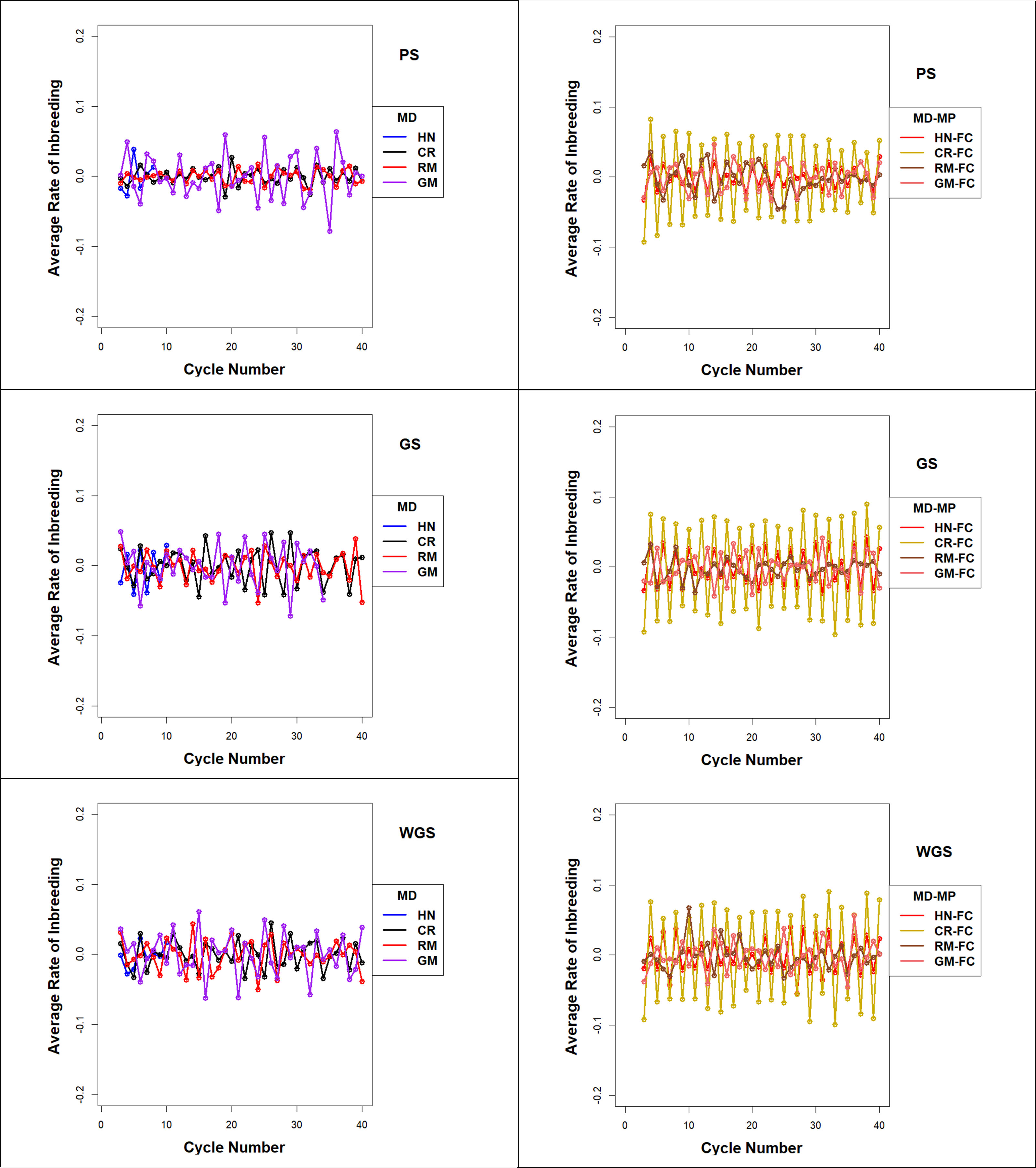

**Supplementary Figure 7.** Average Rate of Inbreeding for Non-isolated (NI) and family island populations (FI) with FC migration policy using PS (top), GS (middle) and WGS (bottom) for the four Mating designs including HN, CR, RM, and GM with top 10% selected fraction. GP models are updated every cycle in GS and WGS, ii) Average Rate of Inbreeding PS (top), GS (middle) and WGS (bottom) with Jannink’s weighting function for the four Mating designs including HN, CR, RM, and GM. GS models are updated every cycle. Selection Methods include PS-Phenotypic Selection, GS- Genomic Selection, and WGS -Weighted Genomic Selection with Jannink weighting function. Mating Designs include HN (Hub Network), CR (Chain rule), RM- Random Mating, GM- Genomic Mating method. Migration policies include bi-directional migrations of two migrants every other cycle involving the Best Island (BI), Random Best (RB), and Fully Connected (FC) islands.

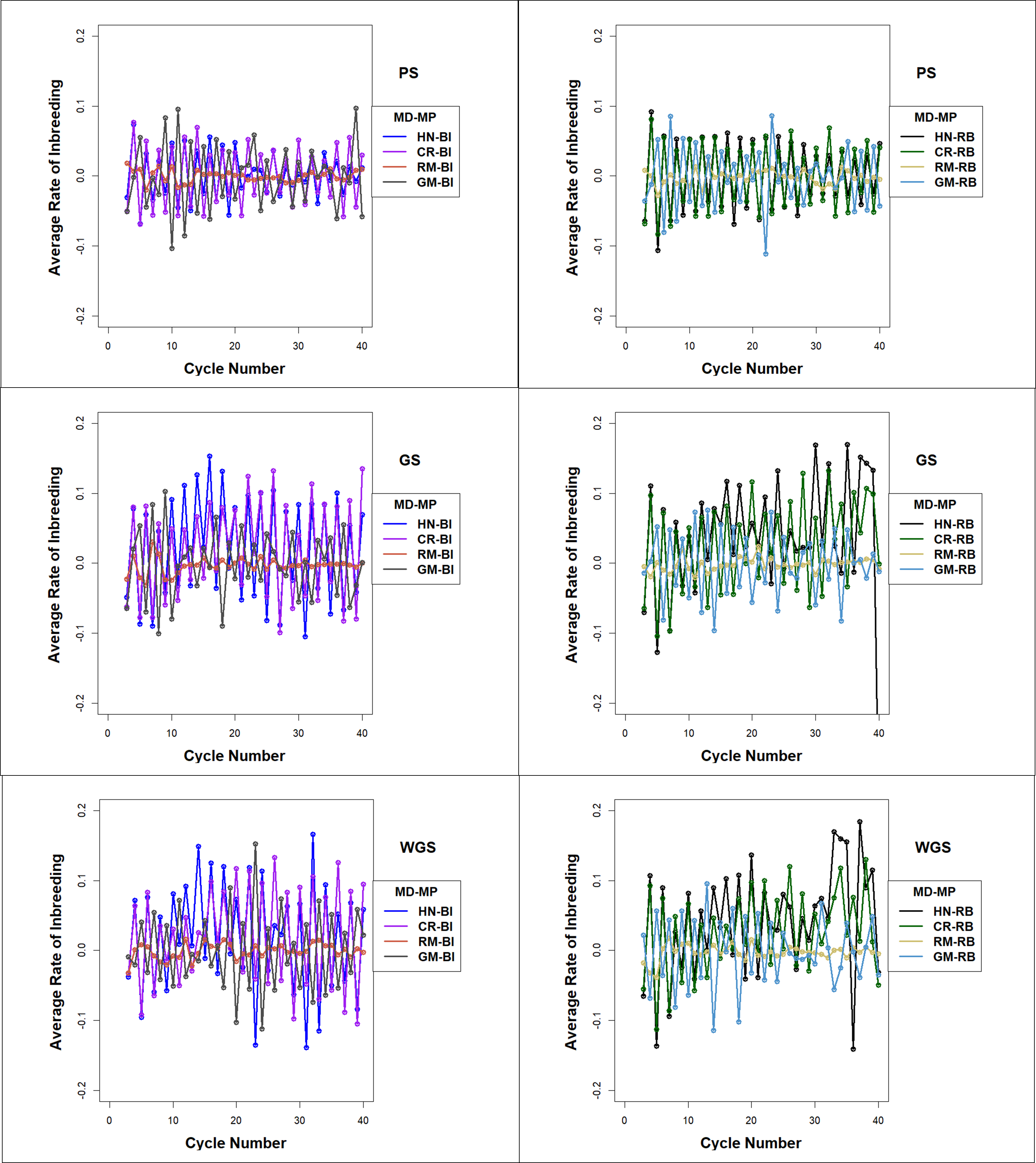

**Supplementary Figure 8.** Average Rate of Inbreeding for island selection with BI and RB migration policies using PS (top), GS (middle) and WGS (bottom) for the four mating designs including HN, CR, RM, and GM with top 10% selected fraction. GP models are updated every cycle in GS and WGS, ii) Average Rate of Inbreeding PS (top), GS (middle) and WGS (bottom) with Jannink’s weighting function for the four Mating designs including HN, CR, RM, and GM. GS models are updated every cycle. Selection Methods: PS-Phenotypic Selection, GS- Genomic Selection, WGS -Weighted Genomic Selection with Jannink weighting function. Mating Designs: HN (Hub Network), CR (Chain rule), RM- Random Mating, GM- Genomic Mating method. Migration policies include bi-directional migrations of two migrants every other cycle involving the Best Island (BI), Random Best (RB), and Fully Connected (FC) islands.

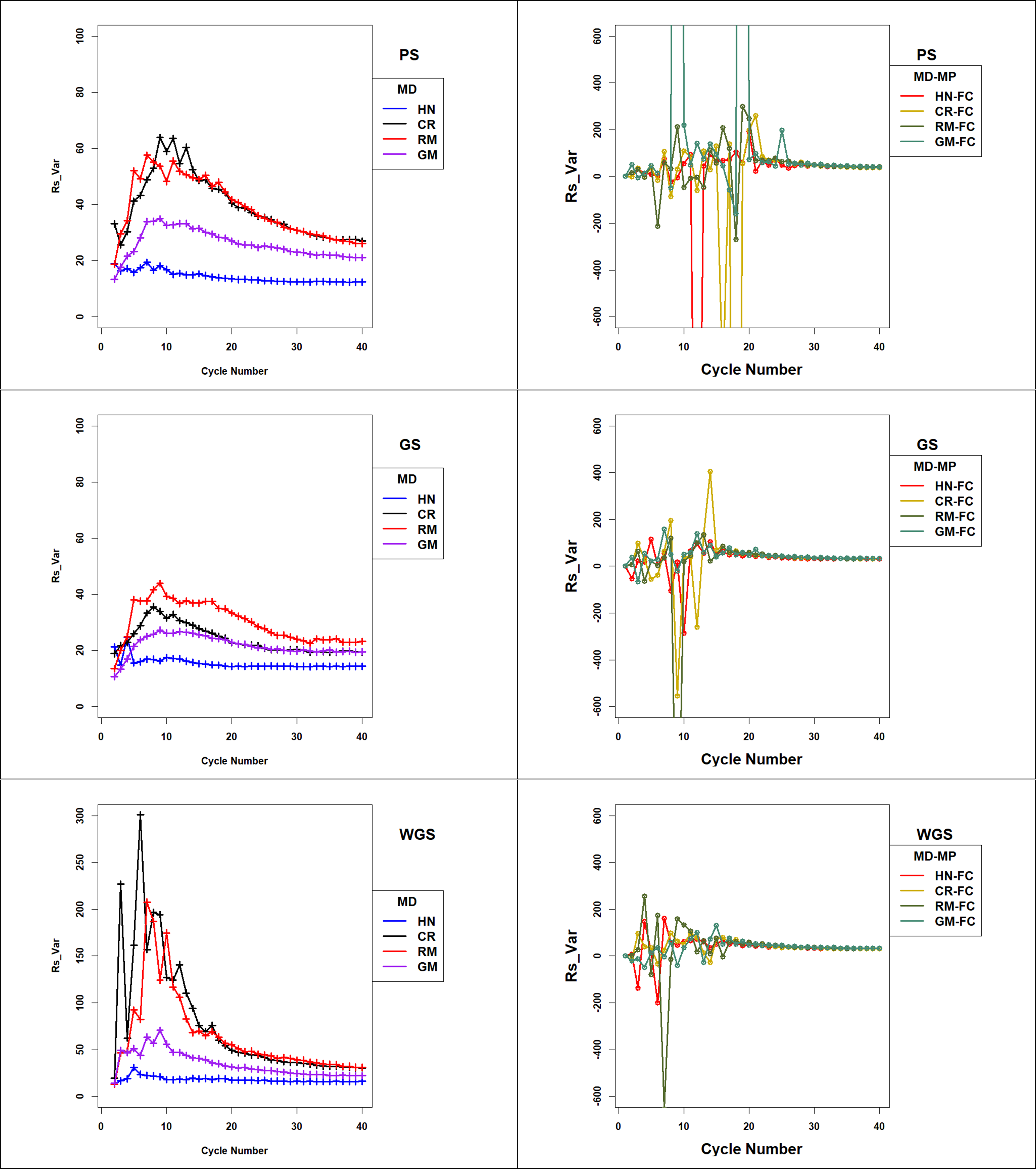

**Supplementary Figure 9 Response Standardized to Change in Genotypic Variance** (**Rs_Var)** across 40 cycles of recurrent selection on non-isolated (left panels) and island (right panels) populations with FC migration policy, using Phenotypic Selection (PS-top row of panels), Genomic Selection (GS-middle row of panels) and Weighted Genomic Selection (WGS-bottom row of panels) and four mating designs: Hub-Network (HN), Chain Rule (CR), Random Mating (RM), and Genomic Mating (GM). Ten percent of lines are selected for crosses using HN, CR, RM and GM mating designs. The genetic architecture in the initial simulated founder lines consisted of 400 additive QTL uniformly distributed throughout the genome and expressed broad sense heritability of 0.7 on an entry mean basis. Genetic variance is standardized to the average genetic variance in founder populations in cycle ‘0’. Average island genetic variance refers to genetic variance within families averaged across 20 families for the four mating designs consisting of HN, CR, RM, and GM. Migration policies include bi-directional migrations of two migrants every other cycle involving the Best Island (BI), Random Best (RB), and Fully Connected (FC) islands.

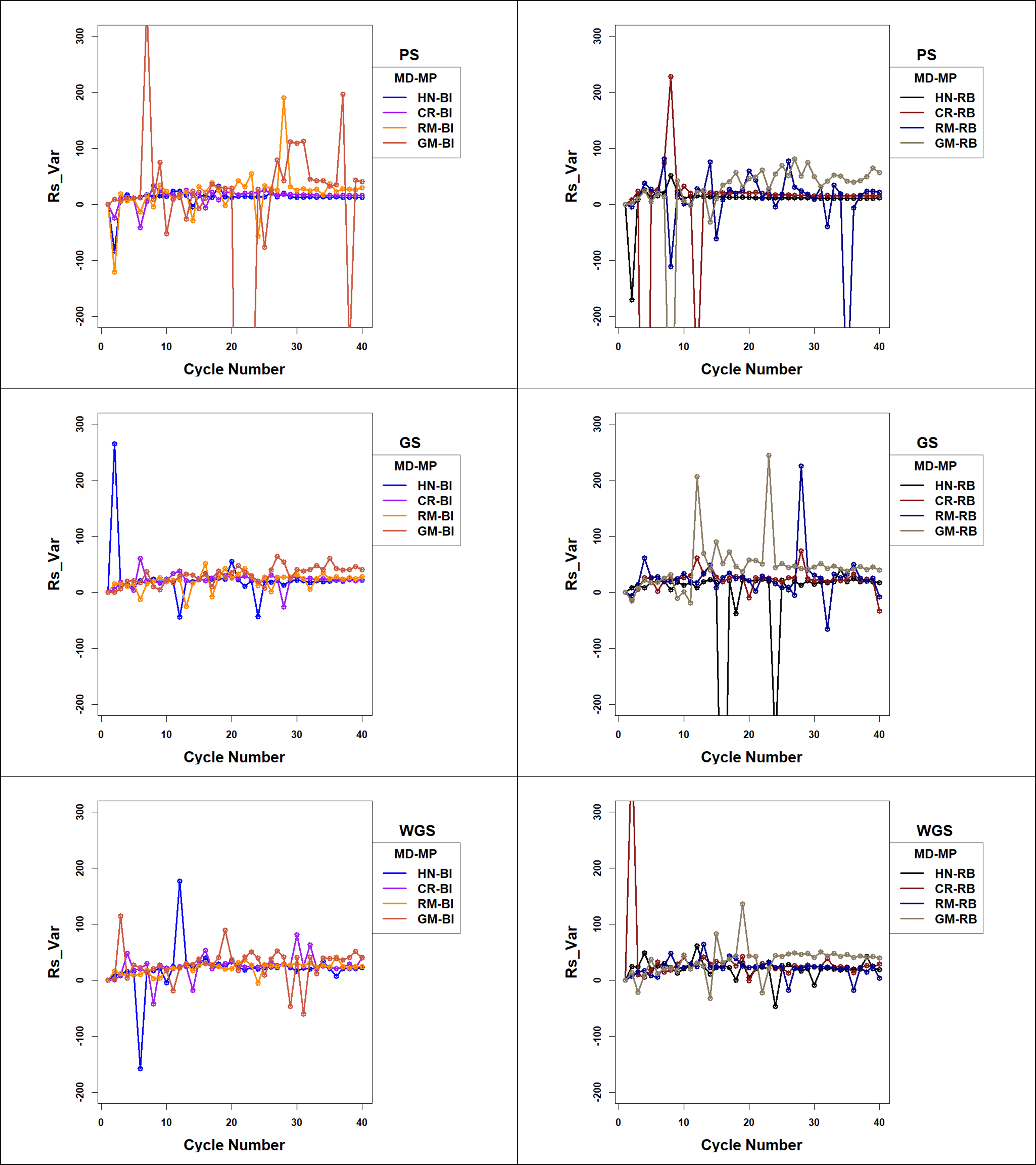

**Supplementary Figure 10 Response Standardized to Change in Genotypic Variance** (**Rs_Var)** across 40 cycles of recurrent selection on island populations with “BI” (left panels) and “RB” migration policies (right panels), using Phenotypic Selection (PS-top row of panels), Genomic Selection (GS-middle row of panels) and Weighted Genomic Selection (WGS-bottom row of panels) and four mating designs: Hub-Network (HN), Chain Rule (CR), Random Mating (RM), and Genomic Mating (GM). Ten percent of lines are selected for crosses using HN, CR, RM and GM mating designs. The genetic architecture in the initial simulated founder lines consisted of 400 additive QTL uniformly distributed throughout the genome and expressed broad sense heritability of 0.7 on an entry mean basis. Genetic variance is standardized to the average genetic variance in founder populations in cycle ‘0’. Average island genetic variance refers to genetic variance within families averaged across 20 families for the four mating designs consisting of HN, CR, RM, and GM. Migration policies include bi-directional migrations of two migrants every other cycle involving the Best Island (BI), Random Best (RB), and Fully Connected (FC) islands.

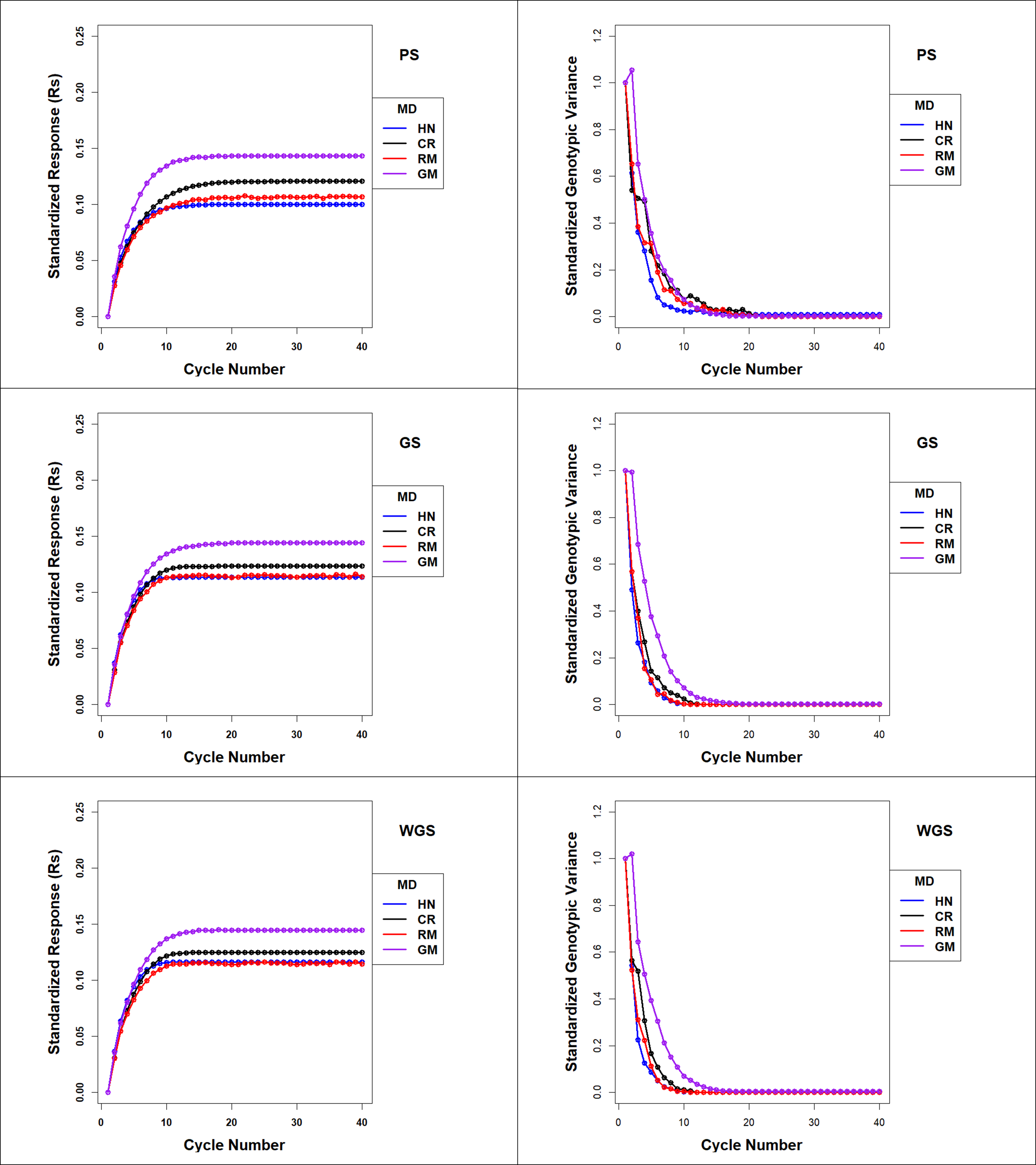

**Supplementary Figure11.** Standardized Response (Rs) and Standardized Genotypic Variance (Sgv) in Discrete selection for PS, GS and WGS s for the four Mating designs including HN, CR, RM, and GM. Standardized genetic response are provided for simulations with 400 simulated QTL responsible for 70% of phenotypic variability. Ten percent of lines are selected from discrete island populations as parental lines for HN, CR, RM and GM designs. Genetic variance is standardized to the average genetic variance in founder populations in cycle ‘0’. Average island genetic variance refers to genetic variance within families averaged across 20 families. GP models are updated every cycle with training data from all prior cycles of selection. Selection Methods: PS-Phenotypic Selection, GS- Genomic Selection, WGS -Weighted Genomic Selection with Jannink weighting function. Mating Design: HN (Hub Network), CR (Chain rule), RM- Random Mating, GM- Genomic Mating method.

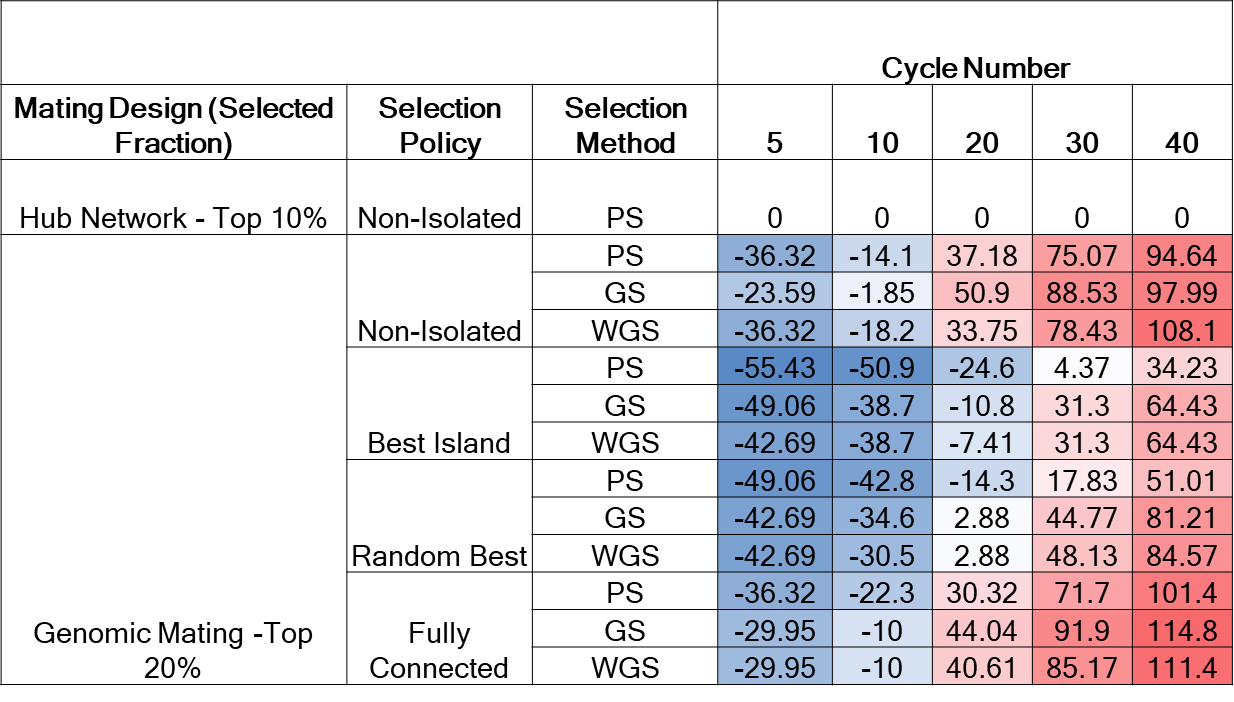

**Supplementary Figure12.** Heat map of relative genetic gain represented as a percentage of genetic response relative to responses in NI-PS-HN in simulations for 400 simulated QTL responsible for 70% of phenotypic variability. Top 10% of the lines are selected for the reference design (NI-PS-HN) for comparisons with responses provided in Supplementary Figure4 and top 20% of the lines are selected as parental lines to be crossed for the GM design in non-isolated and island populations. Selection Methods include PS-Phenotypic Selection, GS- Genomic Selection, and WGS -Weighted Genomic Selection with Jannink weighting function. Mating Designs include HN (Hub Network), GM- Genomic Mating method. Migration policies include bi-directional migrations of two migrants every other cycle involving the Best Island (BI), Random Best (RB), and Fully Connected (FC) topology.
