## Supplemental Table for "Strategies to assure optimal trade-offs among competing objectives for genetic improvement of soybean"

| **Table S1 Weighting Functions for Weighted Genomic Selection** | | | |
| --- | --- | --- | --- |
| **Weighting Function** | **Formula** | **Variables** | **Reference** |
| Jannink Method | 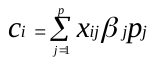 | c_i_ – criterion value for ith individual  x_ij_ – genotype matrix for n- individuals X p-markers  β_j_- effect of marker j  p_j_ – frequency of favorable allele at locus j | Jannink 2010 |
| Dynamic Weighting Method | 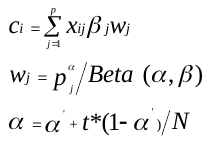 | c_i_ – criterion value for ith individual  x_ij_ – genotype matrix for n-individuals X p-markers  β_j_- effect of marker j  w_j_ – weight at locus j  p_j_ – frequency of favorable allele at locus j  α - shape parameter at generation t for beta distribution  β - shape parameter for beta distribution  t - generation number  N - time horizon | Liu et al. 2015 |
